# Epsin2, a novel target for multiple system atrophy therapy via α-synuclein/FABP7 propagation

**DOI:** 10.1101/2022.06.16.496509

**Authors:** An Cheng, Ichiro Kawahata, Yifei Wang, Wenbin Jia, Tomoki Sekimori, Yi Chen, Nadia Stefanova, David I Finkelstein, Wenbo Ma, Min Chen, Takuya Sasaki, Kohji Fukunaga

## Abstract

Multiple system atrophy (MSA) is a neurodegenerative disease showing accumulation of misfolded α-synuclein and myelin disruption. However, the mechanism how α-synuclein (α-syn) accumulate in MSA brain remains unclear. Here, we identify the protein epsin-2 as a novel target for MSA therapy via controlling α-synuclein accumulation. In MSA mouse model, PLP-hαSyn transgenic mice and FABP7/α-syn hetero-aggregates injected mice, we firstly found that fatty acid-binding protein 7 (FABP7) related to MSA development and formed hetero-aggregates with α-syn, which exhibited stronger toxicity than α-syn aggregates. Furthermore, injected FABP7/α-syn hetero-aggregates in mice selectively accumulated in oligodendrocytes and Purkinje neurons and cause cerebellar dysfunction. By bioinformatic analyses, the protein epsin-2 expresses in both oligodendrocyte and Purkinje cells was found as a potential target to regulate FABP7/α-syn hetero-aggregates propagation via clathrin-dependent endocytosis. The AAV5-dependent epsin-2 knock-down mice exhibited decreased levels of aggregates accumulation in Purkinje neurons and oligodendrocytes as well as performed improved myelin levels and Purkinje neurons in cerebellum and motor functions. Thus, we propose epsin-2 as a novel and therapeutic candidate for MSA.

## INTRODUCTION

Multiple system atrophy (MSA) is an adult-onset, fatal neurodegenerative disease, clinically characterized by progressive autonomic failure, parkinsonism, and signs of cerebellar and pyramidal tract dysfunction ^[1]^. The pathological hallmark of MSA is the abnormal accumulation of misfolded α-synuclein (α-syn), a protein containing 140 amino acids and expressed in neuronal presynaptic terminals ^[2]^, in glial cells, forming the so-called glial cytoplasmic inclusions (GCI). Therefore, MSA is also classified as an α-synucleinopathy together with Parkinson’s disease (PD), dementia and Lewy body disease (DLBD) ^[3]^. However, in contrast to PD or DLBD where α-syn forms neuronal inclusions called Lewy bodies, in MSA, α-syn forms GCI in oligodendrocytes, and seemingly in higher numbers than Lewy bodies in PD and DLBD ^[4]^. Importantly, while higher levels of soluble α-syn have been found in MSA ^[4]^, insoluble α-syn is present at higher levels in PD and DLBD ^[5]^. However, the mechanisms underlying the different expression patterns of α-syn as well as their implications, possibly including different pathological processes and propagation models of α-syn, remain unclear ^[6]^.

In particular, the origin of α-syn composing the GCIs in oligodendrocytes remains unknown. In this sense, the presence of a neuronal protein such as α-syn in oligodendrocytes in MSA has drawn much attention, although several studies have reported no alterations in α-syn mRNA levels in brains of patients with MSA ^[7, 8]^. One hypothesis for the abnormal oligodendroglial expression of α-syn involves cell-to-cell propagation ^[9, 10]^. In this regard, glial cells have been reported to endocytose α-syn from the media and transmit it to neighboring cells, and a direct transfer of neuronal α-syn aggregates has been shown to occur from neurons to glia ^[11]^. Moreover, in rats injected with AAV-hua-syn, a viral vector that induces α-syn expression specifically in neurons, the observed presence of GCIs in oligodendrocytes was caused by α-syn transference from neurons ^[12]^. Importantly, learning about neuronal α-syn transfer to oligodendrocytes contributes to the understanding of how MSA begins and progresses.

In recent years, fatty acid-binding proteins (FABPs), a family of transporter proteins involved in lipid trafficking, have become important factors associated with motor and cognitive functions ^[13-15]^. In dopaminergic neurons, FABP3 regulates dopamine D2L receptor functions ^[16]^ and accelerates α-syn oligomerization in mice treated with 1-methyl-1,2,3,6-tetrahydropiridine (MPTP), which triggers dopaminergic neuron loss ^[15]^. Consistently, we found in a previous study that FABP7, exclusively expressed in glial cells, forms hetero-oligomers and aggregates with α-syn in oligodendrocytes ^[17]^. In addition, previous studies also suggested that FABP7 controls lipid raft formation by regulating the expression of caveolin-1, a protein involved in the endocytic process ^[18]^. Therefore, we hypothesized that FABP7 is involved in the pathological process of MSA and related to the expression pattern and propagation of α-syn in oligodendrocytes.

In this study, we aimed to elucidate the potential role of FABP7 in α-syn accumulation and aggregation in oligodendrocytes and we identified epsin-2 as a potential regulator of α-syn propagation. The obtained results will hopefully contribute to clarifying the mechanism of α-syn aggregation as well as its propagation pattern, which in turn underlies the demyelination and cell loss that occurs during the development of MSA.

## RESULTS

### FABP7 is involved in MSA pathology and α-syn aggregation

In our previous work, we reported that FABP7 formed high-molecular-weight oligomers and aggregates with α-syn in primary oligodendrocytes and glial cells ^[17]^ and argued that the aggregation and accumulation of α-syn in oligodendrocytes were considered the main cause of cell death in MSA. In order to evaluate the role of FABP7 in MSA, we used an *in vivo* genetic model: PLP-hαSyn transgenic mice (PLP-hαSyn mice), in which α-syn is overexpressed in oligodendroglia under the proteolipid protein (PLP) promoter ^[19-21]^ (Fig. 1a). To identify the molecular factors involved in central nervous system (CNS) dysfunction in MSA, a group of selected genes (P ≤0.05) from the RNA sequencing analyses of PLP-hαSyn mouse brain (GSE129531) was measured by GO_KEGG analysis. Consistent with previous work concluding that cerebellar ataxia is the second consensus statement on the MSA diagnosis ^[22]^, we have found a prominent relationship between the MSA condition and the olfactory and spinocerebellar ataxia pathways (Fig. 1b). Particularly in the cerebellum, we found an evident aggregation of hα-syn that was associated with FABP7 (Fig. 1c) in PLP-hαSyn mice. These results suggest a possible interaction between FABP7 and α-syn in MSA pathogenesis. Indeed, in the basal ganglia and cerebellum, we observed co-localization of hαSyn and FABP7 in oligodendrocytes (Fig. 1d). However, the observation that hαSyn also accumulated in Purkinje cell layer of cerebellum mainly in Purkinje cells and basket cells (GABA positive cells) in PLP-hαSyn mice, was unexpected (Fig. 1e). This accumulation might be related to motor dysfunction and Purkinje cell loss (Fig. S1a, b).

**Fig. 1:**
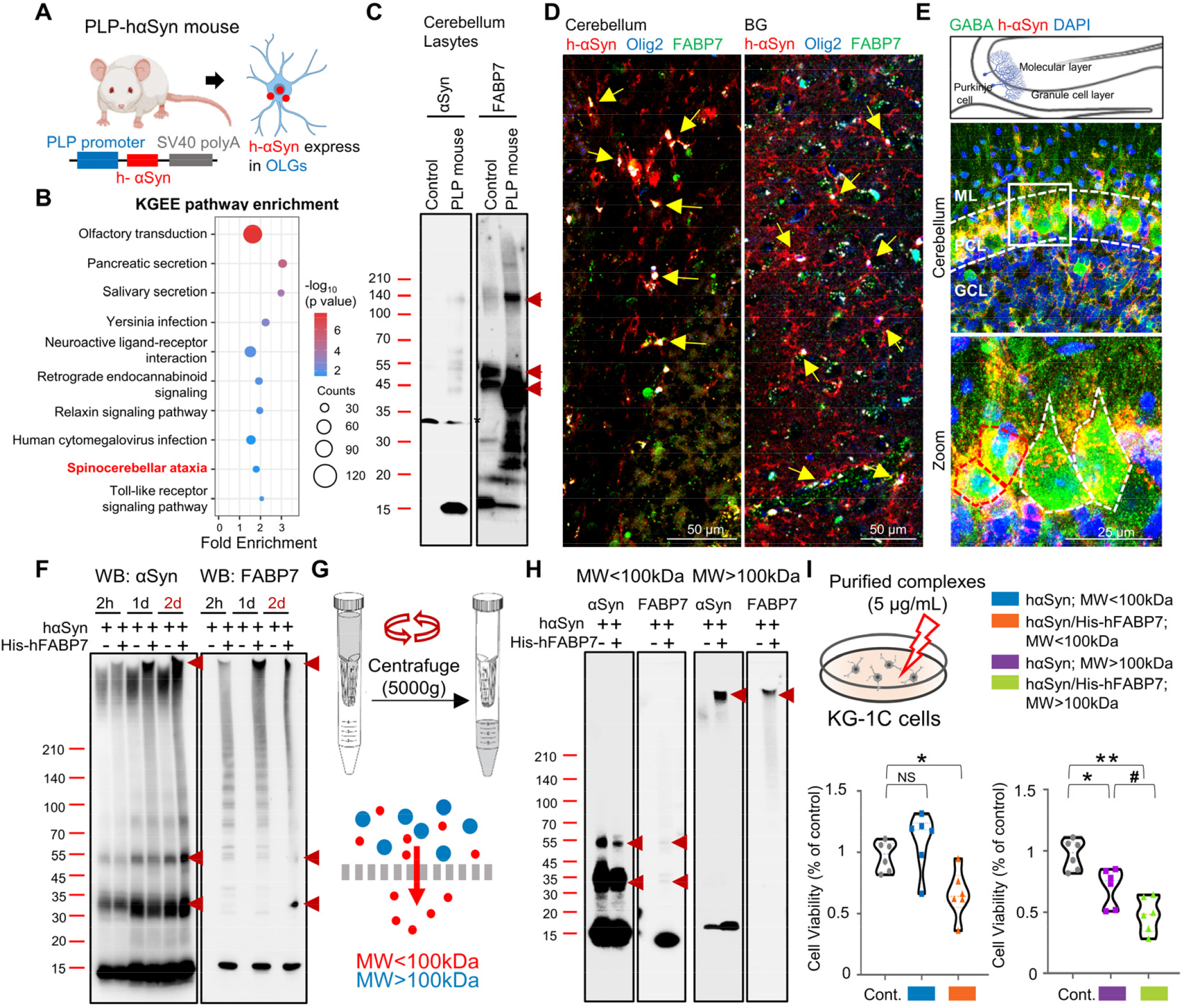
FABP7 related to MSA pathology and participated in αSyn aggregation. **a** Scheme of PLP-hαSyn mice. **b** KEGG analysis of pathways enriched for genes that are differentially expressed in the PLP-hαSyn mouse brain. **c** Western blot analysis of αSyn (left) and FABP7 (right) in the cerebellar extracts of PLP-hαSyn mice. **d** Confocal images show the colocalization of FABP7 (green), hαSyn (red), and Olig2 (blue) in oligodendrocytes in the basial ganglia and cerebellum of PLP-hαSyn mice. **e** Confocal images showing the colocalization of GABA (green), hαSyn (red), and DAPI (blue) in the cerebellum of PLP-hαSyn mice; basket cells (circled by red), Purkinje cells (circled by white). **f** Western blot analysis of αSyn and FABP7, human-αSyn, and human-FABP7 incubated in Tris-HCl (10 mM, pH7.5) for two days. **g** Scheme of the filter used to separate the recombinant proteins. **h** Western blot analysis of αSyn and FABP7. Mixtures were separated into two fractions, below 100 kDa or over 100 kDa. **i** Cell viability analysis of KG-1C cells using the CCK assay. The cells were cultured in separate fractions at a final concentration of 5 μg/ml for 24 h. The data are presented as mean ± standard error of the mean and were obtained using one-way analysis of variance. ^#^P < 0.05, *P < 0.05, and **P < 0.01. αSyn, alpha-synuclein. FABP7, fatty acid-binding protein 7; NS, non-significant; GCL, granule cell layer; PCL, Purkinje cell layer; ML, molecular layer.

We incubated the recombinant proteins hαSyn and His-hfabp7 *in vitro*, in Tris-HCl buffer at 37°C. With longer incubation times, we found an increase in the levels of high molecular weight hαSyn forms. Importantly, in the presence of His-hfabp7, bound hαSyn formed much larger aggregates, particularly after 2 days of incubation (Fig. 1f), however, His-hfabp7 itself could not form high molecular weight aggregates (Fig. S2a, b). Since the presence of FABP7 altered the expression pattern and molecular weight of α-syn aggregates, we thought necessary to identify the differences between the two aggregates, and to determine their toxicity levels. Thus, we isolated the high molecular weight fractions (molecular weight > 100 kDa) using an Ultracel-100 regenerated cellulose membrane (Fig. 1g), and the aggregates were divided into two fractions (molecular weight > 100 kDa or < 100 kDa) (Fig. 1h). To study the differences in toxicity of the two aggregates, we treated KG-1C human oligodendroglial cells with the purified proteins and measured cell viability. Whereas cells treated with hαSyn at low molecular weights (molecular weight < 100 kDa) showed no significant change in cell viability, cells treated with hαSyn/His-hFABP7 at low molecular weights (molecular weight < 100 kDa) showed decreased cell viability (control vs. hαSyn/His-hFABP7 aggregates; MW<100kDa, n=6, p < 0.05, F (2, 15)=8.055). Importantly, cells treated with both hαSyn (Control vs. hαSyn aggregates; MW>100kDa, n=6, p<0.05, F (2, 15)=18.31) and hαSyn/His-hFABP7 (Control vs. hαSyn/His-hFABP7 aggregates; MW>100kDa, n=6, p<0.01) at high molecular weight showed decreased cell viability, although hαSyn/His-hFABP7-treated cells showed a much lower cell viability than hαSyn-treated cells (hαSyn aggregates; MW>100kDa vs hαSyn/His-hFABP7 aggregates; MW>100kDa, n=6, p < 0.05) (Fig. 1i). Therefore, these results indicate that FABP7 not only forms aggregates together with α-syn but also regulates the toxicity of α-syn aggregates.

### FABP7/α-syn hetero-aggregates are selectively taken-up by oligodendrocytes and Purkinje neuron

Our results suggest that FABP7 forms higher molecular weight aggregates with α-syn than with itself and that hetero-aggregates are more toxic. However, it still unclear whether it occurs *in vivo*. Therefore, we further purified hαSyn/His-hFABP7 aggregates, injected them into the lateral ventricles (LV) of wild-type mice (3 μg per mouse) (Fig. 2a) and evaluated their toxicity and propagation *in vivo*. After two months of aggregate injection, we first confirmed the stability of the injected aggregates. Lysates from injected mouse brains were immunoprecipitated and immunoblotted with anti-6X-His and anti-α-syn antibodies; a tight binding of the two proteins was found, indicating that the FABP7/α-syn connection and aggregates were still stable after two months of injection (Fig. 2b). Next, we used whole-brain staining with the 6X-His antibody to measure the distribution of aggregates after injection. We observed that aggregates tended to propagate to the olfactory bulb, cortex, basial ganglia, and cerebellum (Fig. 2c). We then performed a series of behavioral tests to evaluate cognitive function at 4 and 8 weeks post-injection. In the rotarod task test, we observed significant differences in hαSyn/His-hFABP7 aggregate-injected mice at 4 and 8 weeks (4 weeks: n=7, p < 0.05; 8 weeks: n=7, p < 0.05) as well as hαSyn aggregate-injected mice at 8 weeks (8 weeks: n=7, p < 0.05) compared with control mice (Fig. 2d). Similarly, in beam-walking tasks we observed significant differences in hαSyn/His-hFABP7 aggregate-injected mice at 4 and 8 weeks (4 weeks: n=7, p < 0.05; 8 weeks: n=7, p < 0.01) and hαSyn aggregate-injected mice at 8 weeks (8 weeks: n=7, p < 0.05) when compared to control mice. Here, we also observed significant differences between hαSyn/His-hFABP7 aggregate-injected mice and hαSyn aggregate-injected mice at 8 weeks (8 weeks: n=7, p < 0.05) (Fig. 2e). In the Y-maze task, we only observed a significant difference in total arm entry (8 weeks: control vs. hαSyn/His-hFABP7 aggregates; MW>100kDa, n=7, p < 0.01; hαSyn aggregates; MW>100kDa vs hαSyn/His-hFABP7 aggregates; MW> 100kDa, n=7, p < 0.05), but observed no changes in alternation (Fig. 2f, g). These results may indicate that hαSyn/His-hFABP7 aggregates generate a selective toxicity in different brain areas than hαSyn aggregates; however, both aggregate patterns showed a lack of effects on cognitive function.

**Fig. 2:**
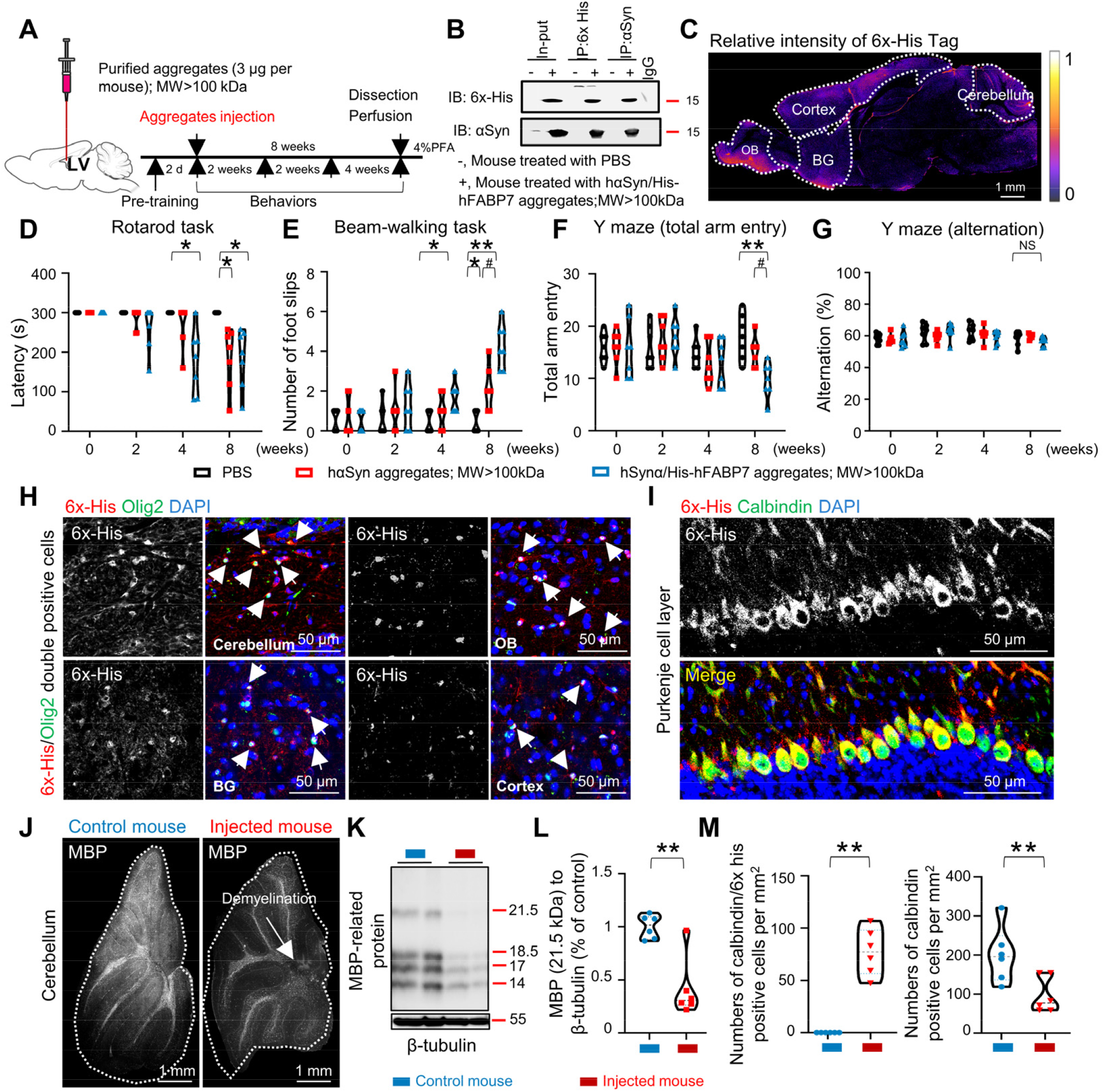
FABP7/α-syn hetero-aggregates suggest motor nerve related toxicity *in vivo*. **a** Scheme of aggregate injection and experimental protocol. Purified aggregates were injected into the lateral ventricles at 3μg per mouse. **b** Co-immunoprecipitation of the 6X His-tag and αSyn in tissue lysates of the olfactory bulb. The extracts were immunoprecipitated (IP) with anti-His or anti-αSyn antibodies. The immunoprecipitates were then immunoblotted (IB) with anti-His or anti-αSyn antibodies. As a control, His and αSyn immunoreactive bands and tissue extracts (input) are shown on the left. **c** Confocal images showing the distribution of aggregates in the whole brain (anti-6X His-tag) after injection for 8 weeks. **d**-**g** Behavioral tasks were carried out at the beginning and after aggregate injection for 2, 4, and 8 weeks. **h** Confocal images showing the colocalization of Olig2 (green), 6X His-tag (red), and FABP7 (blue) in oligodendrocytes. **i** Confocal images show accumulated FABP7/α-syn hetero-aggregates (6x his-tag, red) in Purkinje cells (calbindin, green) **j** Confocal images show decreased MBP levels in FABP7/α-syn hetero-aggregates injected cerebellum. **k** Western blot analysis of MBP and β-tubulin. **l** Quantification of **k** suggested decreased MBP levels in FABP7/α-syn hetero-aggregates injected into the cerebellum. **m** Quantification of calbindin-positive cells and calbindin/6X His-tag double-positive cells, suggesting decreased numbers of calbindin-positive cells and increased numbers of calbindin/6X His-tag double-positive cells. The data are presented as the mean ± standard error of the mean and were obtained using Student’s t-test and two-way analysis of variance. *P < 0.05, **P < 0.01, and ^#^P < 0.05. αSyn, alpha-synuclein; FABP7, fatty acid-binding protein 7; MBP, myelin basic protein; NS, nonsignificant. BG, basal ganglia; OB, olfactory bulb.

After the injection of hαSyn/His-hFABP7 aggregates for 2 months, the aggregates mainly propagated to four regions: cerebellum, olfactory bulb, basal ganglia, and cortex. In these regions, we found that aggregates were selectively taken up by oligodendrocytes (Fig. 2h) as well as accumulated in the purkinje cell layers (Fig. 2i). Even though relatively low levels of aggregates were taken up by oligodentrotyes in the cerebellum, this may still account for by the significant decrease in MBP in the white matter of the cerebellum in the hαSyn/His-hFABP7 aggregate-injected mice (control mouse vs. injected mouse, n=6, p < 0.01) (Fig. 2j, k, l). On the other hand, the Purkinje neurons also significantly decreased in number (control mouse vs. injected mouse, n=6, p < 0.01) accompanied by accumulation of hαSyn/His-hFABP7 aggregates (control mouse vs. injected mouse, n=6, p < 0.01) compared with control mice (Fig. 2m). Therefore, these results indicate that FABP7/α-syn hetero-aggregates show distinct patterns of α-syn aggregation, suggesting higher toxicity towards oligodendrocytes and Purkinje neurons, possibly due to selective uptake.

### Scanning for potential receptors which regulating FABP7/α-syn hetero-aggregates propagation

To date, we have already identified the role of FABP7 in α-syn aggregate formation and in their higher toxicity towards oligodendrocytes and Purkinje neurons, possibly due to a differential mechanism by which these cells selectively take up FABP7/α-syn heteroaggregates. This is reminiscent of a specific receptor that selectively modulates FABP7/α-syn hetero-aggregate propagation and may be involved in the MSA pathological process. Interestingly, in a previous work, FABP7 was also reported to control lipid raft function through the regulation of caveolin-1 expression in astrocytes ^[18]^. Thus, considering the important role of FABP7 in aggregation, we performed a transcriptomic profile analysis of the datasets to identify candidate genes. We identified 20 genes that were significantly changed in all datasets (Fig. 3a). However, single-cell sequencing data from the Sequence Read Archive (SRA) (GSM3449587)^[23, 24]^ suggested that only six genes (ENP2, FOXO3, KRT4, SLC27A3, SLC1A2, and RAB27B) were expressed in both oligodendrocytes and Purkinje neurons (Fig. 3b, c; Fig. S5). Next, as a result of protein-protein correlation analysis, we found the gene FABP7 and EPN2, which relates to cell endocytosis ^[25]^ are strongly correlated (Fig. 3c). Interestingly, the mRNA expression of ENP2 was consistently downregulated in the whole blood of patients with MSA with FABP7, based on RNA-seq analysis data (GSE34287)^[26]^ (Fig. 3d).

**Fig. 3:**
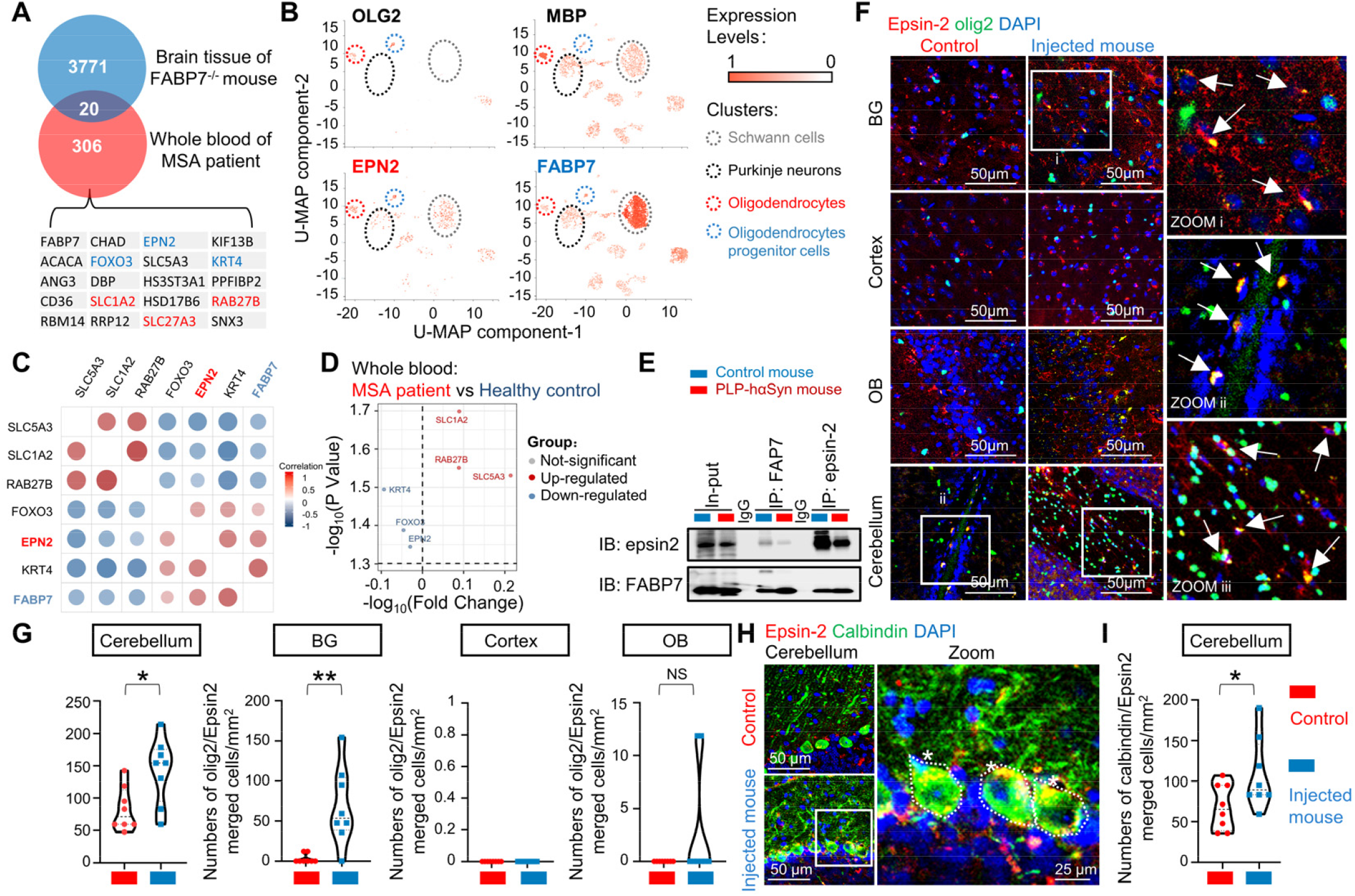
Scanning for potential receptors. **a** Overlapping Venn diagram of transcriptomic differential genes in two candidate gene databases. **b** UMAP dimensional reduction visualizations of 5240 cells from the olfactory bulb (mouse strainC57BL/6NJ). The color represents the normalized log expression of the selected genes. Each point represents a single cell. Original data were obtained from the Gene Expression Omnibus (GEO; GSM3449587), and sequencing data were handled using PanglaoDB (https://panglaodb.se/). These results suggest that six genes (ENP2, FOXO3, KRT4, SLC27A3, SLC1A2, and RAB27B) were expressed in both oligodendrocytes and Purkinje neurons. **c** Protein-protein correlation analysis for the six selected genes, revealing two specific interactions, marked with a star symbol. **d** Volcano plot of the log2 fold-change (n=3) on the X-axis and log^10^ of the p-value derived using two-way ANOVA. Each point represents an individual mRNA expression. Colored points represent significantly changed mRNA, and gray points represent non-significant mRNA (p < 0.05, |log2 FC| ≥ 0.01). **e** Co-immunoprecipitation of Epsin-2 and FABP7 in tissue lysates of the cerebellum. Extracts were immunoprecipitated (IP) with anti-Epsin-2 or anti-FABP7 antibodies. The immunoprecipitates were then immunoblotted (IB) with an anti-Epsin-2 antibody or anti-FABP7 antibody. **f** Confocal images showing the distribution of epsin-2 (red) in oligodendrocytes (olig2, green), and DAPI (blue). **g** Quantification of G suggested increased levels of olig2 and epsin-2 merged areas in the cerebellum and basal ganglia. **h** Confocal images showing the distribution of epsin-2 (red) in Purkinje neurons (olig2, green), and DAPI (blue). **i** Quantification of **h** suggested increased levels of epsin-2 in Purkinje neurons. The data are presented as the mean ± standard error of the mean and were obtained using the Student’s t-test. **P < 0.01. αSyn, alpha-synuclein; FABP7, fatty acid-binding protein 7; MBP, myelin basic protein; NS, nonsignificant; BG, basal ganglia; OB, olfactory bulb.

To further identify whether the gene ENP2 is involved in FABP7/α-syn hetero-aggregate propagation, we first immunoprecipitated both the epsin-2 protein, encoded by the ENP2 gene, and FABP7. We could observe that epsin-2 binds to FABP7 in both control and PLP-hαSyn mice (Fig. 3e). This result corroborated our hypothesis that epsin-2 is involved in regulating FABP7/α-syn heteroaggregate propagation. We then verified the cellular distribution of epsin-2 in the brain by immunostaining. We found that epsin-2 is abundantly expressed in oligodendrocytes (Fig. 3f) and Purkinje neurons (Fig. 3h) but barely in neurons and astrocytes (Fig. S4a, b) in both control and injected mice. Importantly, we found increased epsin-2 levels in oligodendrocytes in cerebellum (control mouse vs. injected mouse, n=8, p < 0.05), and BG (control mouse vs. injected mouse, n=8, p < 0.01) of aggregates injected mouse. Consistently, epsin-2 levels in Purkinje neurons are upregulated in aggregates injected mice (Fig. 3h, i) (control mouse vs. injected mouse, n=8, p < 0.01). The results may indicate that epsin-2 levels were consistent with those of FABP7/α-syn hetero-aggregates in injected mice. Therefore, these results indicate a close link between epsin-2 and FABP7 in aggregate formation; in particular, epsin-2 may play a role in FABP7/α-syn hetero-aggregate propagation.

### Knock-down of epsin-2 decreased FABP7/α-syn hetero-aggregates uptake and cerebellum dysfunction

When we injected mice with FABP7/α-syn hetero-aggregates into the lateral ventricles of the brain, they exhibited motor dysfunction after four weeks, and loss of Purkinje neurons or demyelination mainly in the cerebellum. Thus, we then performed cerebellar injection with FABP7/α-syn hetero-aggregates (2 μg per mouse). To identify the role of epsin-2 in FABP7/α-syn heteroaggregate propagation, we injected mice with a viral vector (pAAV-mEPN2 shRNA-EGFP-U6) that knocked down epsin-2 levels. The viral vectors were injected 2 weeks before aggregate injection, and motor functions were measured by beam-walking and rotarod tasks once a week (Fig. S7a, b). We observed significant differences in the beam walking test at 1-, 2-, and 3-weeks post-injection in mice injected with hαSyn/His-hFABP 7 aggregates compared to control mice (1 week: Control vs. aggregates injection, n=5, p < 0.01; 2 weeks: Control vs. aggregates injection, n=5, p < 0.01; 3 weeks: Control vs. aggregates injection, n=5, p < 0.01). The rotarod task also resulted in significant differences at 1, 2, and 3 weeks in mice injected with hαSyn/His-hFABP 7 aggregates compared to control mice (1 week: Control vs. aggregates injection, n=5, p < 0.05; 2 week: Control vs. aggregates injection, n=5, p < 0.01; 3 week: Control vs. aggregates injection, n=5, p < 0.01) (Fig. S7c). However, we did not observe significant differences following viral infection in neither of the motor tests, beam-walking and rotarod, at 3 weeks (Fig. S7c). This lack of effects in motor functions may be due to the limited area of viral infection. In the infected area, we found decreased levels of hαSyn/His-hFABP7 aggregates in oligodendrocytes (Aggregates Injection mouse vs. Aggregates + AAV Injection mouse, n=5, p < 0.01) and Purkinje neurons (Aggregates Injection mouse vs. Aggregates + AAV Injection mouse, n=5, p < 0.01) as well as decreased levels of episn-2 (Fig. S7d, e). Consistently, in oligodendrocyte precursor cell (OPCs) primary culture, the virus-infected cells also showed significantly decreased levels of hαSyn/His-hFABP7 aggregate uptake (Fig. S6).

To get higher and wider knock-down efficiency of epsin-2, we next injected viral vector into the lateral ventricles of the brain 2 weeks before hαSyn/His-hFABP7 aggregates injection (Fig. 4a). Followed the viral vector spread to the whole brain (Fig. 4b), we found decreased episn-2 and accumulated aggregates levels in both Purkinje cells and oligodendrocytes in cerebellum (Fig. 4c, d) (Purkinje cells: aggregates injection vs. aggregates +AAV injection, n=8, p < 0.01; oligodendrocytes: aggregates injection vs. aggregates +AAV injection, n=8, p < 0.05). Furthermore, we also observed improved moto functions in viral vector injected mice. At this time, we observed significant differences in the beam walking test at 8-weeks in mice injected with viral vector compared to hαSyn/His-hFABP7 aggregates injected mice (8 week: aggregates injection vs. aggregates +AAV injection, n=5, p < 0.05) (Fig. 4e). The rotarod task also resulted in significant differences at 8 weeks in mice injected with viral vector compared to hαSyn/His-hFABP7 aggregates injected mice (8 week: aggregates injection vs. aggregates +AAV injection, n=5, p < 0.05) (Fig. 4f).

**Fig. 4:**
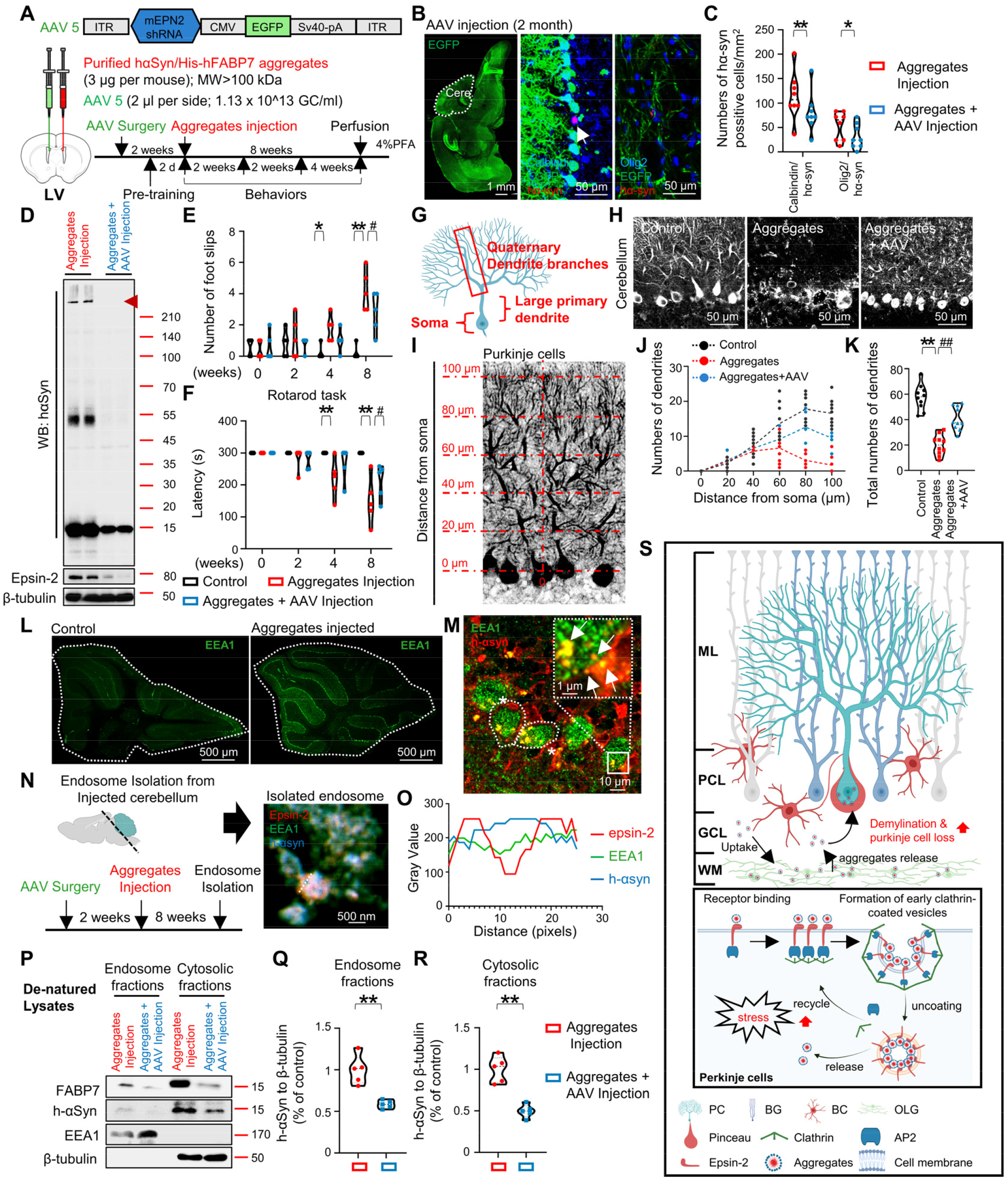
Knock-down of epsin-2 decreased FABP7/α-syn hetero-aggregates dependent toxicity. **a** Scheme of aggregates, AAV injection, and experimental protocol. **b** Confocal images showing the distribution of AAV (green) in Purkinje neurons (calbindin, red), oligodendrocytes (olig2, red) and DAPI (blue) in the cerebellum. **c** Quantification of B, suggested decreased levels of hαSyn/His-hFABP7 aggregates uptake in purkinje neurons and oligodendrocytes in AAV injected mice. **d** Western blot analysis of αSyn, epsin-2, and β-tubulin, suggested deceased levels of αSyn in episn-2 knock-down mouse. **e, f** Behavioral tasks were carried out at the beginning and after aggregates or AAV injection for 2, 4, and 8 weeks. **g** Scheme of Purkinje neuron morphology. **h** Confocal images showing the calbindin in green, DAPI in blue. **i, j** Verify Purkinje neuron dendrites by length. **k** Quantification of total dendrites of Purkinje neuron, suggested improved Purkinje neuron morphology by episn-2 knock-down. **l** Confocal images showing EEA1 (green) expression in cerebellum. **m** Confocal images showing colocalization of EEA1 and αSyn in FABP7/α-syn hetero-aggregates injected mouse. **n** Scheme of endosome isolation from Injected cerebellum (left) and confocal images showing colocalization of EEA1, epsin-2, and αSyn. **o** Fluorescence intensity change in specific area. **p** Western blot analysis of FABP7, h-αSyn, EEA1, and β-tubulin in endosome fractions and cytosolic fractions. **q, r** Quantification of **p** suggested decreased levels of αSyn in both endosome and cytosol fractions isolated from episn-2 knock-down mouse. **s** Schematic representation of the pathways of αSyn accumulation and propagation in cerebellum. Injected αSyn aggregates were selectively take up by oligodendrocytes and Purkinje cells via episn-2-dependent endocytosis thereby causing demyelination and Purkinje cell loss in cerebellum. The data are presented as the mean ± standard error of the mean and were obtained using Student’s t-test and two-way analysis of variance *P < 0.05, **P < 0.01, ^#^P < 0.05, and ^##^P < αSyn, alpha-synuclein; FABP7, fatty acid-binding protein 7; MBP, myelin basic protein; NS, nonsignificant; PC, Purkinje cell; PCL, Purkinje cell layer; BG, Bergmann glia; GCL, granule cell layer; BC, basket cell; OLG, oligodendrocytes; AP2, adaptor protein 2; ML, molecular layer; WM, white mater.

On the other hand, when we focus on the Purkinje neurons morphology, which closely related cerebellum functions (Fig. 4g). We found improved morphology (Fig. 4h) in length (Fig. 4i, j) and numbers (Fig. 4k) (8 week: aggregates injection vs. aggregates +AAV injection, n=5, p < 0.01) of dendrites of Purkinje neurons in viral vector injected mice. Thus, these results show that epsin-2 regulates the uptake of hαSyn/His-hFABP7 aggregates in oligodendrocytes and Purkinje neurons thereby affecting cerebellar function.

### Epsin-2 regulates FABP7/α-syn hetero-aggregates propagation via endocytosis

Epsin-2 is reported as an endocytic adapter protein that contain binding sites for clathrin^[27]^ and involved in clathrin-dependent endocytosis. Thus, we hypothesis that the large size of FABP7/α-syn hetero-aggregates were taken up into cells via this pathway. When we stained cerebellum with endosome biomarker: EEA1, we found up-regulated levels of EEA1 in Purkinje cell layer of aggregates injected mice (Fig. 4l) and colocalization of EEA1 and hα-syn in very small size areas (100-500 nm) (Fig. 4m). It may demonstrate activated endocytosis response to aggregates injection.

We next isolate endosomes from aggregates injected cerebellum (Fig. 4n). By confocal microscopy, we found hα-syn inside endosome and coated by epsin-2 (Fig. 4o). Moreover, in Epsin-2 knock-down mice we also found decreased α-syn levels in both endosome fractions (Fig. 4p, q) (8 week: aggregates injection vs. aggregates +AAV injection, n=5, p < 0.01) and cytosolic fractions (Fig. 4p, r) (8 week: aggregates injection vs. aggregates +AAV injection, n=5, p < 0.01). These results demonstrated that epsin-2 regulated FABP7/α-syn hetero-aggregates propagation via endocytosis.

## DISCUSSION

According to the clinical features, MSA is divided into two main types: the parkinsonian type (MSA-P) with poor levodopa-responsive parkinsonian syndrome and the cerebellar type (MSA-C) with cerebellar ataxia syndrome ^[28]^. The degree of morphological changes in the cerebellum is more pronounced in MSA-C ^[29]^ including Purkinje neuron dysfunction, such as an increase in torpedoes in Purkinje axonal morphology and reduction in spines of Purkinje dendritic arbor; these changes are in turn implicated in cerebellar dysfunction ^[30]^. Moreover, the loss of Purkinje neurons and increased levels of GCIs in the cerebellum of MSA patients may be due to demyelination and disease progression ^[31]^. Consistently, in our study, we also observed an obvious accumulation of α-syn in Purkinje neurons and oligodendrocytes in hαSyn/His-hFABP7 aggregate-injected mice. In MSA patients, there is strong evidence that α-syn accumulation in oligodendrocytes results in loss of myelin and oligodendroglial dysfunction ^[32]^ within the pons and cerebellum, which is in turn related to the presence of GCI and disease duration ^[33]^. Loss of Purkinje neurons is another important cerebellar feature associated with MSA-C ^[29, 34]^. Thus, Purkinje neuron loss seems to be a secondary consequence of α-syn accumulation and demyelination. We hypothesize that, besides the neuroinflammation in MSA-C ^[35]^ induced by demyelination, the direct propagation of α-syn from oligodendrocytes to Purkinje neurons is a key factor for Purkinje neuron degeneration.

Since the etiology of various α-syn-related diseases, such as PD and MSA, remain unclear, it is possible that distinct conformations of α-syn aggregates result in different clinical outcomes ^[36-38]^. In contrast to the PD brain, the MSA brain contains soluble activity, and the α-syn form of the MSA brain induces distinct inclusion morphologies ^[39]^. Consistently, in contrast to the insoluble α-syn amyloid fibrils in the PD brain, in our work, the FABP7/α-syn heteroaggregates constructed by 2-days-incubation of recombinant protein were soluble. Due to the limitations of our experimental conditions, we cannot conclude that the pattern of α-syn aggregation in the MSA brain is the same as that of FABP7/α-syn hetero-aggregates in PLP-hαSyn mice; however, FABP7/α-syn hetero-aggregate propagation is completely different from α-syn amyloid fibrils in wild-type mice. In previous studies, α-syn preformed fibril (PFFs) treatment led to the accumulation of α-syn in dopamine neurons and selective decreases in synaptic proteins ^[40]^ as well as motor deficits ^[41]^. In contrast, in the present study, FABP7/α-syn hetero-aggregates only propagated to oligodendrocytes and Purkinje neurons in FABP7/α-syn hetero-aggregate-injected mice. Evidently, the propagation of distinct α-syn strains into different regions in the CNS should involve a specific receptor.

The α-syn protein is intracellular and cytosolic. For a successful propagation of α-syn to additional brain regions, the aggregates must bind to the cell surface and entry into the cytoplasm. A previous study indicated that dopamine D2 long receptors were critical for α-syn uptake ^[42]^ towards dopamine neurons. In this study, we found that epsin-2 is a potential receptor that regulates the propagation of aggregates in MSA. Epsin-2 is an endocytic adapter protein that contains an N-terminal ENTH domain along with unfolded central and C-terminal regions that contain binding sites for clathrin ^[27]^. Clathrin does not bind to the plasma membrane directly, and epsin-2 is a ubiquitous component of endocytic clathrin-coated pits that interacts with lipids on the plasma membrane ^[43]^. Consistently, by confocal microscopy, we found the α-syn was wrapped in epsin-2 and formed endosomes inside cells. Furthermore, the knock-down of epsin-2 also decreased α-syn levels of oligodendrocytes and Purkinje neurons. In RNA sequencing analysis, we found that the EPN2 gene strongly correlated with FABP7 and was significantly changed in the whole blood of patients with MSA. By immunostaining and immunoprecipitation analysis, we also demonstrated that epsin-2 regulated FABP7/α-syn hetero-aggregate propagation towards oligodendrocytes and Purkinje neurons, and the virus infected-epsin-2 knock-down mice exhibit improved motor functions, which related with improved Purkinje morphology and myelin levels. These results demonstrated that α-syn propagation was regulated by epsin-2 expression especially in oligodendrocytes and Purkinje neurons via clathrin-dependent endocytosis.

Here still some limitations in our study that we only identified our hypothesis in preclinical studies using MSA model mice, we lack clinical evidence to support our hypothesis that FABP7 is involved in α-syn aggregation of GCI formation directly and epsin-2 also regulated α-syn aggregates accumulation in MSA brain. On the other hand, even though the epsin-2 knock-down mice which infected with virus exhibit significant improvement of motor functions, it is a pity that, so far, none of the specific episn-2 inhibitors can be obtained from the market. Thus, we are planning to clarify the mechanism and binding site for epsin-2 to regulate α-syn propagation and furthermore develop specific ligands for epsin-2 inhibition in our future work.

In this study, we verified the vital role of FABP7 in α-syn aggregation in MSA and identified a special propagation pattern of FABP7/α-syn hetero-aggregates that selectively propagate to oligodendrocytes and Purkinje neurons. Importantly, epsin-2 is a novel receptor that regulates FABP7/α-syn hetero-aggregates propagation via endocytosis. Additionally, the knock-down of epsin-2 successfully attenuated Purkinje cell dysfunctions and demyelination in cerebellum. Overall, epsin-2 should be considered as a potential therapeutic target for the development of inhibitors that overcome MSA clinical symptoms.

## MATERIALS AND METHODS

### Key resources table

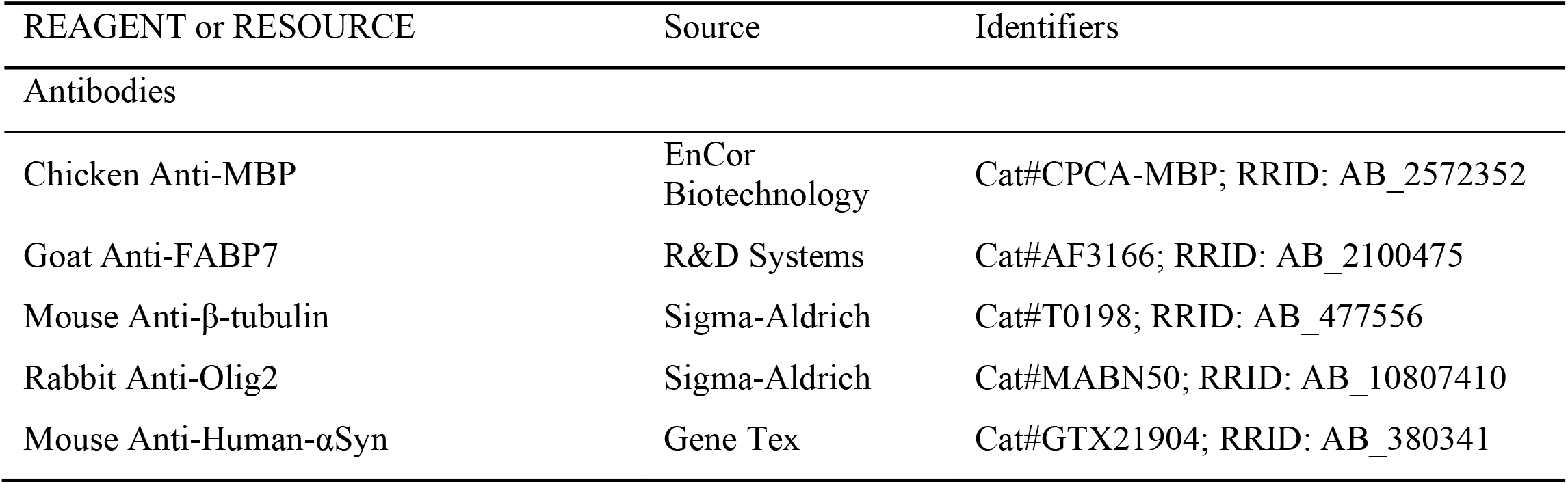
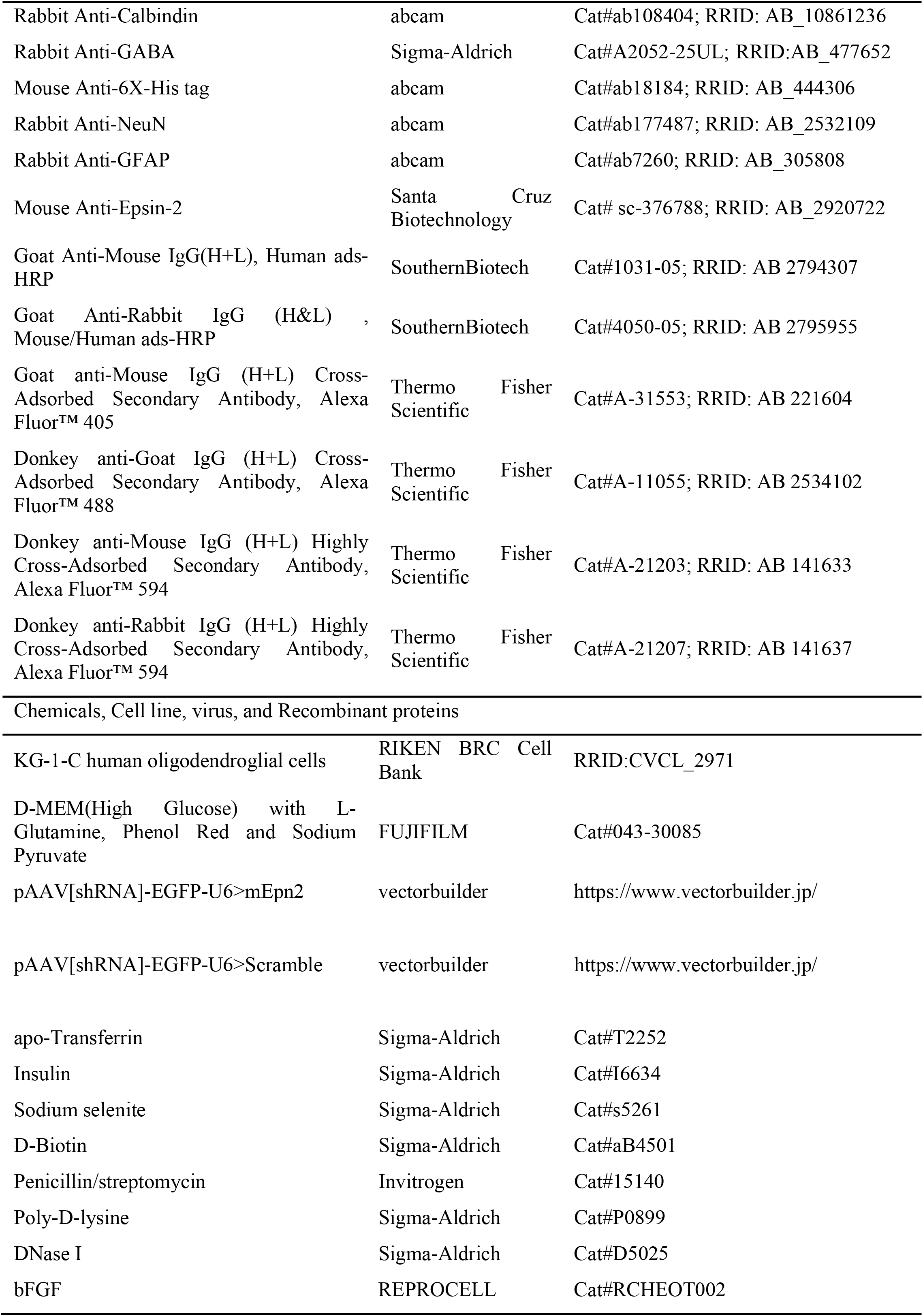
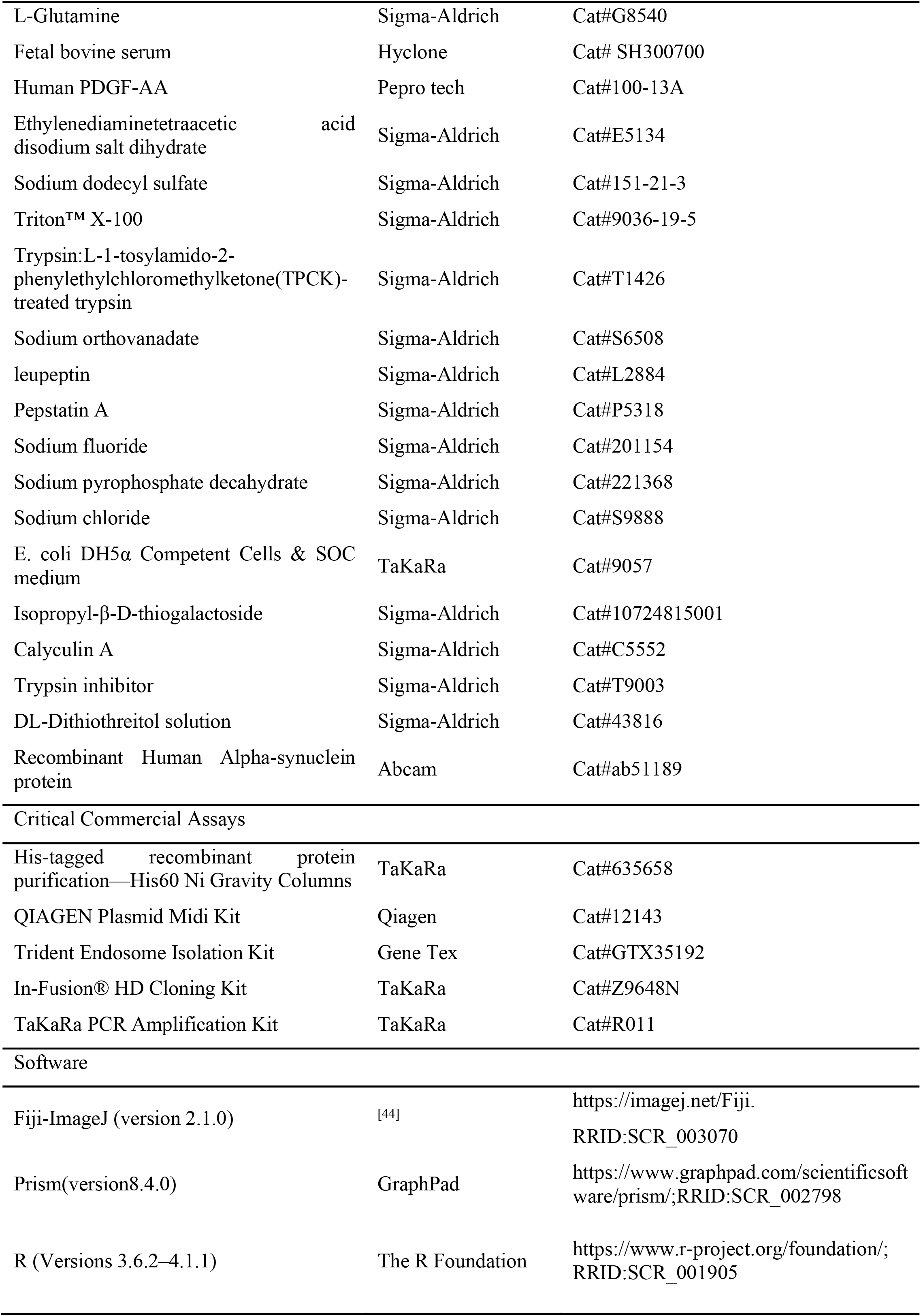

#### Animals

Eight-week-old male C57BMF/6J mice (Clea Japan, Inc., Tokyo, Japan) and PLP-hαSyn transgenic mice were maintained in polypropylene cages (temperature: 23±2°C; humidity: 55±5%; lights on between 9 a.m. and 9 p.m.). Animals had unlimited access to food and water. We used only male mice in this study to minimize the effect of sex hormones such as estrogen. All animal experiments were reviewed and approved by the Institutional Animal Care and Use Committee of Tohoku University Environmental and Safety Committee (2019PhLM0-021 and 2019PhA-024).

Male and female PLP-αsynuclein mice overexpressing human wild-type α-synuclein under the control of the oligodendrocyte-specific proteolipid protein promoter (PLP-αsyn mice) on a C57Bl/6 background were used for all experiments ^[45]^. The mice were housed in a temperature-controlled, pathogen-free environment under a 12-h light/dark cycle at the animal facility of The Florey Institute of Neuroscience and Mental Health and allowed access to standard laboratory chow and water ad libitum. All the experiments were approved by the Florey Institute Animal Ethics Committee (#13-117 and #18-006).

All animal experiments were performed in accordance with animal use guidelines and ethical approval. In each group, the mice were treated and randomly measured, and there were no confounders. Isoflurane was used for anesthesia during the experiment. For euthanasia, the mice were intraperitoneally injected with an overdose of pentobarbital.

#### Primary oligodendrocyte precursor cells (OPCs) culture

The method for primary OPCs culture has been described previously ^[46]^. Briefly, P1-2 mouse pups were decapitated, followed by isolation of the cortex from the brain and digestion in a solution (13.6 mL PBS, 0·8 mL DNase I stock solution [0·2 mg/mL], and 0·6 mL of a trypsin stock solution [0·25%]). The tissues were sliced into approximately 1 mm^3^ pieces using a sterilized razor blade, centrifuged at 100 × *g* for 5 min, and resuspended in DMEM20S medium (DMEM, 4 mM L-glutamine, 1 mM sodium pyruvate, 20% fetal bovine serum [FBS], 50 U/mL penicillin, and 50 μg/mL streptomycin). The tissue suspension was strained using a 70 μm nylon cell strainer, seeded in poly L-lysine-coated tissue culture flasks, and cultured in DMEM20S for 2 weeks (mixed culture). After that The flasks were shaken for 1 h at 200 rpm at 37 °C to remove microglial cells and for an additional 20 h to detach OPCs, and then seeded on poly-D-lysine-coated plates and cultured in OPC medium (DMEM, 4 mM L-glutamine, 1 mM sodium pyruvate, 0.1% BSA, 50 μg/mL Apo-transferrin, 5 μg/mL insulin, 30 nM sodium selenite, 10 nM D-biotin, 10 nM hydrocortisone 10 ng/mL PDGF-AA, and 10 ng/mL bFGF).

#### Preparation of protein

FABP7 with an N-terminal 6X His-tag (His-hFABP7) was expressed in E. coli DH5α competent cells using pET28a derived vector containing coding genes constructed using the In-Fusion® HD Cloning Kit. is. The hFABP7 gene was inserted into pET28a using either the restriction sites NcoI and XhoI (hFABP7), or NdeI and XhoI (His-hFABP7). Expression vectors were confirmed by sequencing before their use for protein expression. For protein purification, after transformed cells were grown to OD_600_=0.6∼0.8, cells were additionally incubated with 1 mM isopropyl-β-D-thiogalactoside (IPTG, 1mM) for 15h at 37 °C. The cells were centrifuged (5000 rpm, 10min, 4 °C) and resuspended in Tris-HCl buffer (50mM Tris-HCl, pH7.4, in a 10:- (w/v) ratio). Next, the resuspended cells were sonicated, and centrifugated (15000rpm, 10min, 4 °C); the collected supernatant was brought to 80% saturation by solid ammonium sulfate. The supernatant containing His-hFABP7 was resolubilized in Tris-HCl buffer (50mM Tris-HCl, pH7.4). For His-hFABP7 extraction, His-tagged recombinant protein purification (His60 Ni gravity columns) was performed following the manufacturer’s instructions. After column development, we pooled the fractions containing His-hFABP7 and added a 0.4-fold volume of diisopropyl ether: n-butanol (3:2) solution. The mixture was incubated at 4 °C with gentle agitation (20 rpm) for three cycles at 30 min intervals. The solution was desalted by sequential dialysis (5 mM, 3 mM, and 1 mM NH4HCO3) and finally lyophilized to obtain the delipidized, lyophilized His-hFABP7 protein.

#### Aggregates construction

Purified His-hFABP7 (0.2 mg / ml), and hαSyn (0.2 mg / ml) obtained from Abcam were incubated in Tris-HCl buffer (150 mM NaCl, 10 mM Tris, pH 7.5) at 37 °C for 2h, 24h, and 2 days. The binding of FABP7 to αSyn was confirmed by western blotting. To separate aggregate fractions, the incubated protein mixture was collected and centrifuged using centrifugal filters Utracel-100K (Millipore) at 500 × g for 10 min. The filter was washed with PBS and centrifuged again at 500 × g for 10 min. Repeat it three times, the last residue on the filter is the aggregate with the molecular weight of over 100 kDa. Aggregates were desalted by sequential dialysis (5, 3, and 1 mM NH4HCO3) and finally lyophilized.

#### Endosome isolation

The endosomes from mouse brain were isolated using Trident Endosome Isolation Kit (GTX35192) according to the manufacturer’s instructions.

#### Cell culture

KG-1C human oligodendroglial cells (RRID: CVCL_2971) were obtained from RIKEN BRC Cell Bank (Tsukuba, Japan). Short-tandem repeat (STR) profiling of the cell lines was performed using JCRB. Cells were cultured in Dulbecco’s modified Eagle’s medium (DMEM) with 10% FBS) and penicillin/streptomycin (100 U/100 μg/mL) at 37°C under 5% CO2.

#### Stereotactic injection

For intracerebroventricular injections, the same titer (1.13×10^13^ GC/ml) and equal amounts (2 μL per side) of viral particles or constructed aggregates (3 μL per mouse) were injected into the lateral ventricle at the following coordinates (anterior—0.7 mm; lateral±1; depth—2.5 mm relative to the bregma) through a Hamilton syringe (Hamilton Company, Reno, NV, USA). For direct cerebellar injection, the same titer (1.13×10^13^ GC/ml) and equal amounts (1 μL per mouse) of viral particles or constructed aggregates (1 μL per mouse) were injected into the cerebellum directly at the following coordinates (anterior—6 mm; lateral 0 mm; depth—2.8 mm relative to the bregma) through a Hamilton syringe (Hamilton Company, Reno, NV, USA).

#### Protein extraction from brain tissue

Frozen tissues were subjected to lysis buffer containing 0.5% Triton-X100, 4 mM ethylene glycol, 50 mM Tris-HCl (pH 7.4), 10 mM ethylenediaminetetraacetic acid, 1 mM sodium orthovanadate, 50 mM sodium fluoride, 40 mM sodium pyrophosphate decahydrate, 0.15 M sodium chloride, 50 μg / mL leupeptin, 25 μg / mL peptatin A, 50 μg / mL trypsin inhibitor, 100 nM cariculin A, 1 mM dithiothreitol. The tissues were then homogenized and sonicated. After centrifugation (15000 rpm, 10 min, 4 °C), the supernatant protein was collected, and the concentration was normalized using Bradford’s assay. Then, the samples were mixed with 6 × Laemmli sample buffer containing no β-mercaptoethanol and treated at 100 °C for approximately 3 min.

#### SDS-PAGE

Extracts (30 μg) were separated by SDS-polyacrylamide gel electrophoresis (SDS-PAGE) using a ready-made gel (Cosmo Bio Co., Ltd.) and transferred to a polyvinylidene fluoride membrane in buffer containing 3.03 g/L Tris, 14.41 g/L glycine for 2 h. The membrane was then blocked with 5% defatted milk powder in TTBS solution containing 50 mM Tris–HCl (pH7.5), 150 mM NaCl, and 0.1% Tween 20 for 1 h. Membranes were incubated overnight with primary antibodies at 4 °C. After washing three times with TTBS (10 min each time), the membrane was incubated with TTBS TTBS-diluted secondary antibody for 2 h at room temperature. The membrane was then incubated using an ECL detection system (Amersham Biosciences, NJ, USA), and intensity quantification was performed using Image Gauge software version 3.41 (Fuji Film, Tokyo, Japan).

#### Slice and histochemistry

The mice were anesthetized with isoflurane and perfused with PBS, followed by 4% PFA. The brain was fixed overnight with 4% PFA and sliced using a cryostat (Leica) (Twenty-micron brain sections). Immunofluorescence staining was performed as previously described^[14]^. The slice was permeated for 5 min using 0.1% Triton X-100 in PBS and then washed three times (10 min each time) and blocked with PBS / 1% BSA for 1 h. The primary antibodies were incubated overnight in blocking solution at 4 °C. After washing thrice with PBS and 1% BSA, a fluorescent secondary antibody was added. Finally, immunofluorescence images were analyzed using a confocal laser-scanning microscope (DMi8; Leica, Wetzlar, Germany).

#### Cell viability assay

Cell viability was measured using a cell counting kit (CCK) (Dojindo), according to the manufacturer’s instructions. This assay is based on the clearance of the water-soluble tetrazolium salt, WST-8. It is reduced by intracellular dehydrogenase to produce a yellow formazan pigment. This dye is soluble in the tissue culture medium. The absorbance in living cells was measured at 400 nm and 450 nm.

#### Y-maze task

Spatial memory was evaluated using a Y-maze task. A Y-shaped maze with three arms (50 cm long, 7.5 cm high, 10.5 cm wide) connected at an angle of 120 °was placed on the desk for use. The mouse was placed at the end of one arm and allowed to explore the maze freely for 8 min, and the positions of the arms to which the mouse moved were recorded in the selected order. The number of times the mouse moved to each arm within the specified time was counted and taken as the total number of arm selections. Based on this, we investigated the combination of three different arms selected in succession, and we were able to recognize this number as the number of alternating actions. In the quantitative calculation, the number of alternation actions is divided by the total number of arm selections minus two, and the value obtained by multiplying the value by 100 is used to obtain the alternation action rate (alterations (%)), which is used for voluntary alternation actions.

#### Rotarod task

A device consisting of iron rods (3 cm in diameter and 30 cm in length) was machined to prevent slipping. In the test, the mouse was placed on a rotating rod at 30 rpm and the time to fall was recorded for up to 300 s.

#### Beam walking task

A (155 mm x 160 mm x 5 mm) “goal box” was placed at the top of a rectangular beam (length:870 mm x width:5 mm) and both ends of the beam were fixed at 500 and 315 mm from the floor. For training, the mice were placed 10, 30, 50, and 80 cm from the goal and walked to the goal box. In the test, the mouse was placed 80 cm from the goal box, and the number of missed steps required to reach the goal box within 60 s was recorded.

#### Bioinformatics

The original RNA sequencing data of PLP-hαSyn mice and whole blood of MSA patients were obtained from the Gene Expression Omnibus (GEO; GSE129531, https://www.ncbi.nlm.nih.gov/geo/query/acc.cgi?acc=GSE129531; GSE34287^[26]^, https://www.ncbi.nlm.nih.gov/geo/query/acc.cgi), and (GSM3449587^[23, 24]^, respectively. The data were processed using RStudio software. The original data for single-cell sequencing were obtained from the Sequence Read Archive (SRA) (GSM3449587)^[23, 24]^ (https://www.ncbi.nlm.nih.gov/geo/query/acc.cgi?acc=GSE121891) and were measured using PanglaoDB^[47]^ (https://panglaodb.se/#google_vignette).

#### Statistics

Data were analyzed using GraphPad Prism 8 (GraphPad Software, Inc., La Jolla, CA, USA) and are expressed as mean ± standard error of the mean (SEM). The normality assumption was examined using the Shapiro-Wilk test, and the equal variance assumption was examined using the Brown-Forsythe test in GraphPad Prism 8. For data that passed these assumptions, significant differences were determined using Student’s t-test for two-group comparisons (Fig. 2l; Fig. 2n; Fig. S2b ; Fig. 3h; Fig. 4e; Fig. S4c; Fig. S6c) and by one-way analysis of variance (ANOVA) or two-way analysis of variance (ANOVA) for other multigroup comparisons, followed by Tukey’s multiple comparisons test. Statistical significance was set at p < 0·05.

## List of Supplementary Materials

Fig. S1 to S7

Table S1 to S2

Full-size scans of western blots

## Acknowledgments

We thank the Uehara Memorial Foundation, Japan Science and Technology Agency (JST) (grant number JPMJCR21P1), and Japan Agency for Medical Research and Development (AMED) (grant numbers JP20dm0107071; 22ym0126095h0001) for their financial support.

## Author contributions

An Cheng, developed the concept, planned and performed experiments, analyzed data, and original draft writing; Ichiro Kawahata, analyzed experiments; Yifei Wang, recombinant protein purification; Wenbin Jia, methodology and mouse behaviors; Tomoki Sekimori, cell culture; Yi Chen, methodology; Nadia Stefanova, manuscript review; David I Finkelstein, animals; Wenbo Ma, methodology; Min Chen, methodology; Takuya Sasaki and Kohji Fukunaga, supervision, project administration, funding, and manuscript review and editing. All authors critically reviewed and approved the final version of the manuscript. A. Cheng and K. Fukunaga verified the data and had the final responsibility for the decision to submit for publication.

## Data and materials availability

The materials used in this study are available commercially. The data supporting the findings of this study are available in the article and supplementary material. The data in Fig. 1b, and Fig. 3a, c, and d were based on publicly available data from the Gene Expression Omnibus (GEO; GSE129531, https://www.ncbi.nlm.nih.gov/geo/query/acc.cgi?acc=GSE129531; GSE34287^[26]^, https://www.ncbi.nlm.nih.gov/geo/query/acc.cgi). The data in Fig. 3b and Fig. S5 were based on publicly available data from the Gene Expression Omnibus (GSM3449587^[23, 24]^, https://www.ncbi.nlm.nih.gov/geo/query/acc.cgi?acc=GSE121891) and measured by PanglaoDB^[47]^ (https://panglaodb.se/#google_vignette). The schematic diagrams in this work were drawn by Biorender (https://biorender.com/) with a publication license. The readers are welcome to contact the corresponding author for the raw data used in this study.

## Supplementary Materials

**Figure S1.**
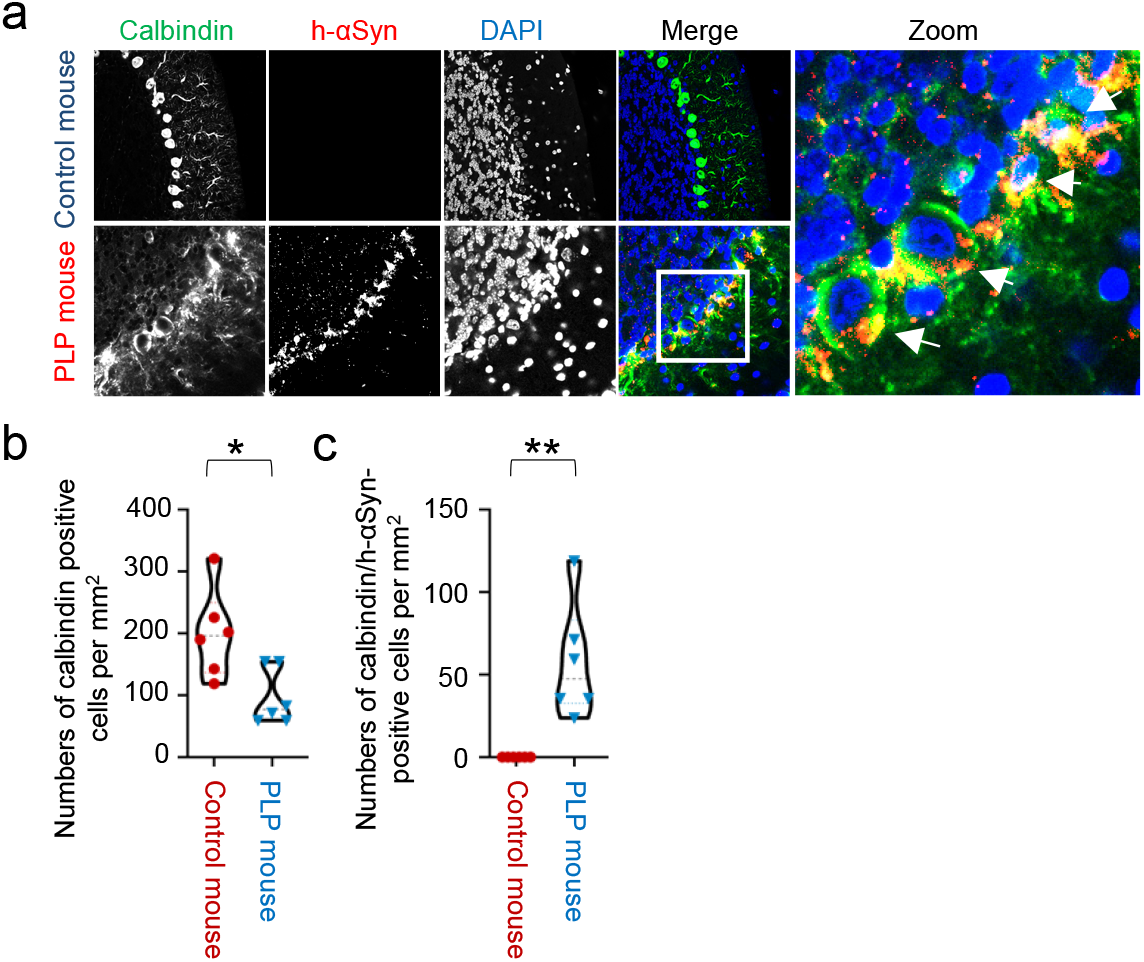
Purkinje cells also revealed MSA pathology association. a, Confocal images show h-αSyn in red, calbindin in green, DAPI in blue. h-αSyn localized in purkinje cells in PLP mouse. b, c Quantification of calbindin positive cells, of calbindin/h-αSyn-positive cells In PLP mouse the numbers of purkinje cells significantly decreased compared with control group and the calbindin/h-αSyn-positive cells increased significantly in PLP mouse. The data are presented as the mean ± the standard error of the mean, were obtained using Unpaired Student’s t-test. *P < 0.05 ; **P < 0.01. αSyn, alpha-synuclein.

**Figure S2.**
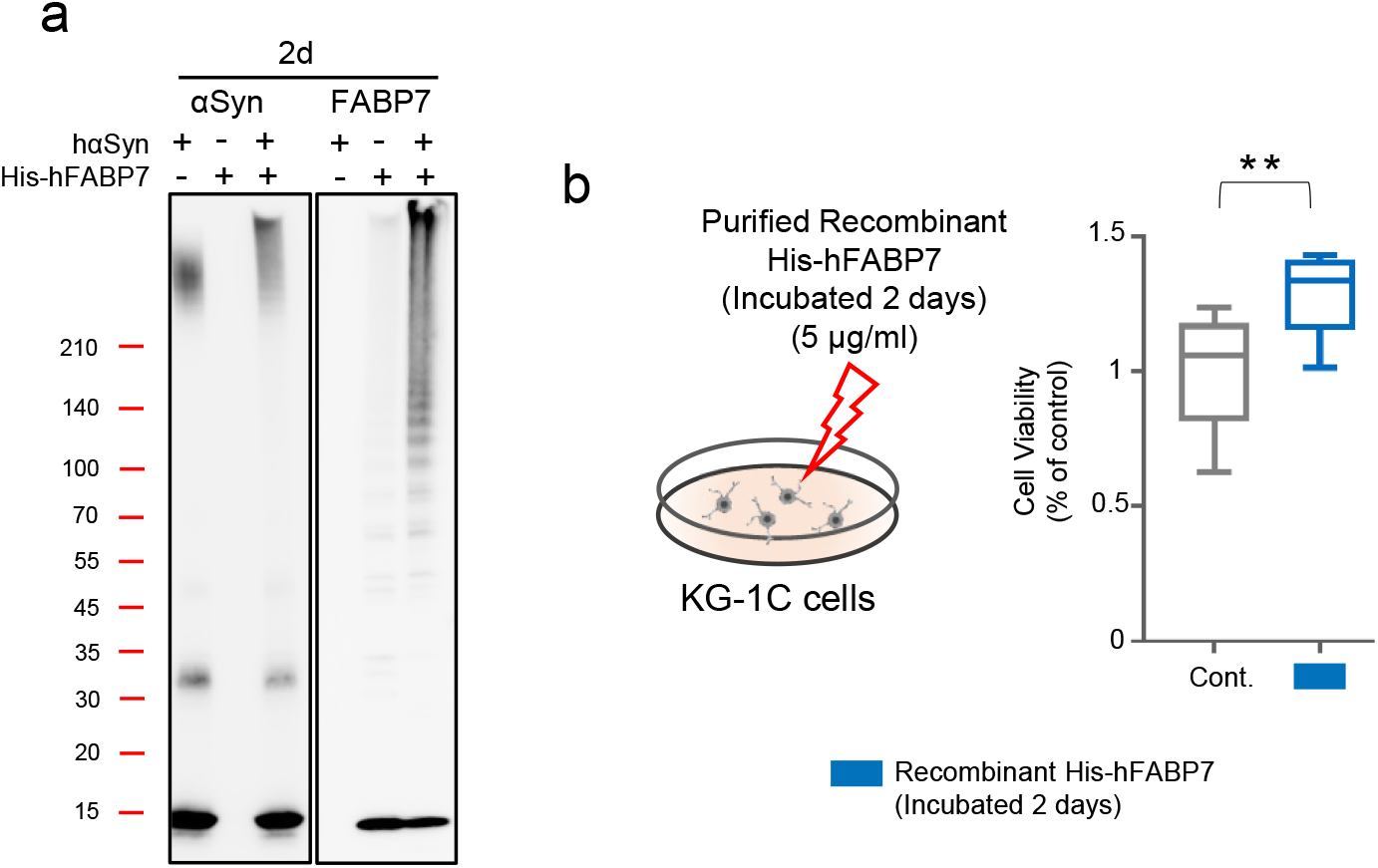
Recombinant His-hFABP7 suggests no obvious toxicity towards oligodendrocytes. a, Western blot analysis of αSyn and FABP7, human-αSyn and human-FABP7 were incubated in Tris-HCl (10 mM, pH=7.5) for two days. Incubation of His-hfabp7 only were not able to trigger high molecular weight aggregates. b, Cell viability analysis of KG-1C cells, based on a CCK assay. Cells cultured treated with separated fractions at a final concentrations of 5 μg/ml, for 24 h. The data are presented as the mean ± the standard error of the mean, were obtained using Unpaired Student’s t-test. **P < 0.01. αSyn, alpha-synuclein. CCK, cell counting kit-8; FABP7, fatty acid binding protein 7.

**Figure S3.**
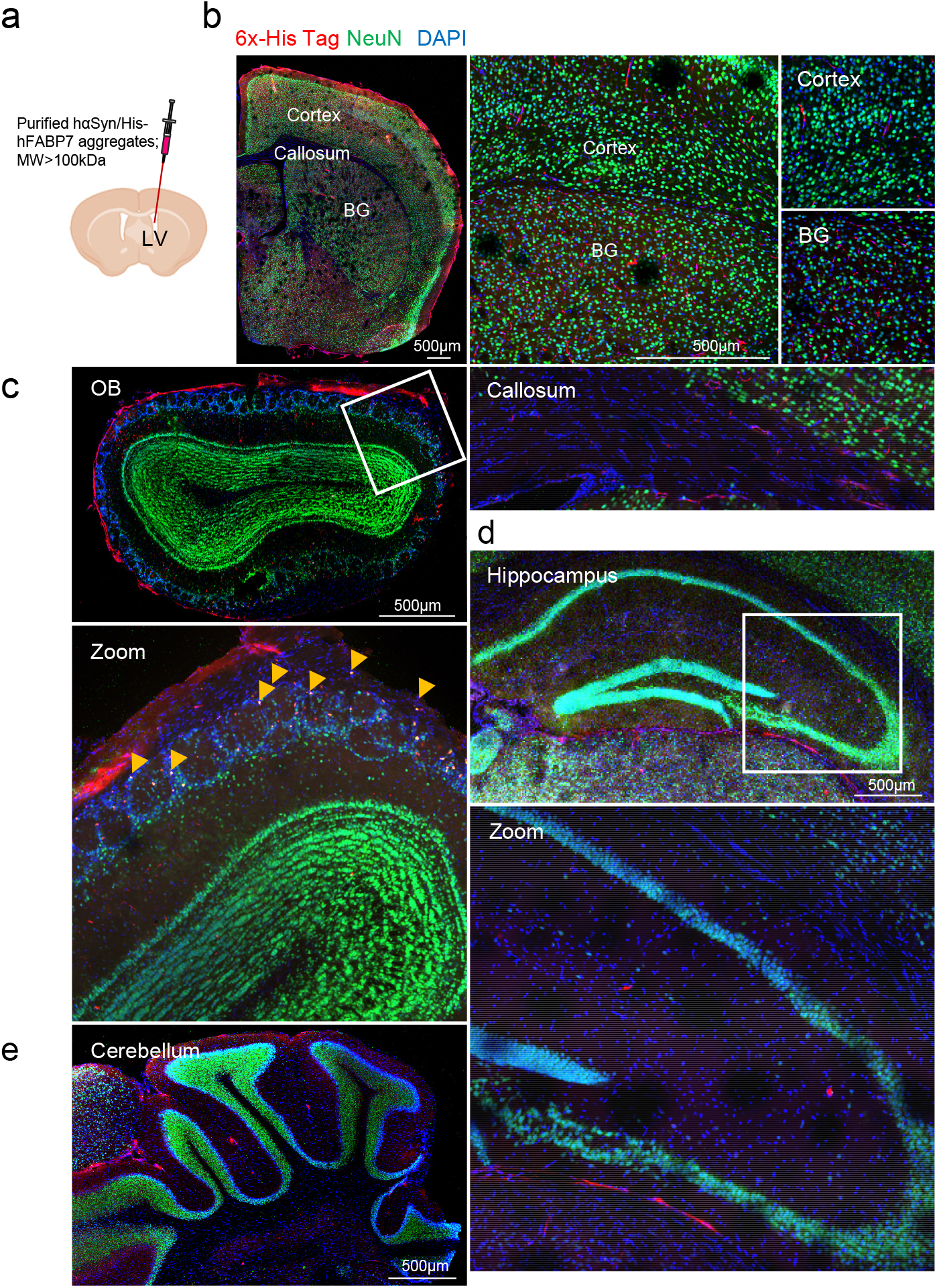
High molecular weight hαSyn/His-hFABP7 aggregates are not selectively taken up by neurons. a, Scheme of aggregates injection, αSyn/FABP7 aggregates were injected into Lateral Ventricles for 3μg per mouse. b, c, d, e, Confocal images show His-tagged aggregates in red, NeuN in green, DAPI in blue. hαSyn/His-hFABP7 aggregates are not selectively taken up by neurons in cortex, BG, hippocampus and cerebellum. But a small number of aggregates were taken up by neurons in OB. BG, basal ganglia; OB, olfactory bulb.

**Figure S4.**
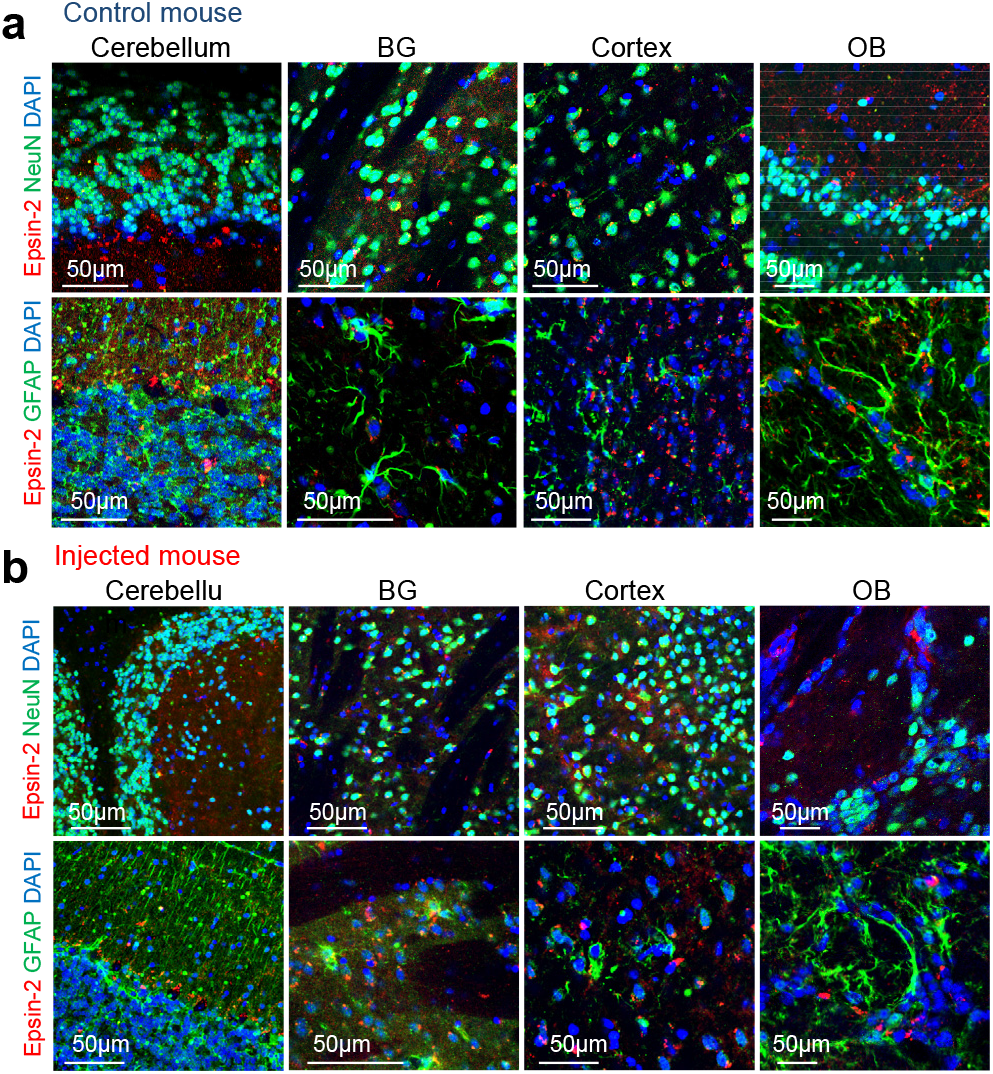
Expression of epsin-2 in neurons and astrocytes. a, b Confocal images show epsin-2 in red, NeuN or GFAP in green, DAPI in blue. Epsin-2 does not express in astrocytes but some part of neurons or in BG and cortex of control mouse and hαSyn/His-hFABP7 aggregates injected mouse. BG, basal ganglia.

**Figure S5.**
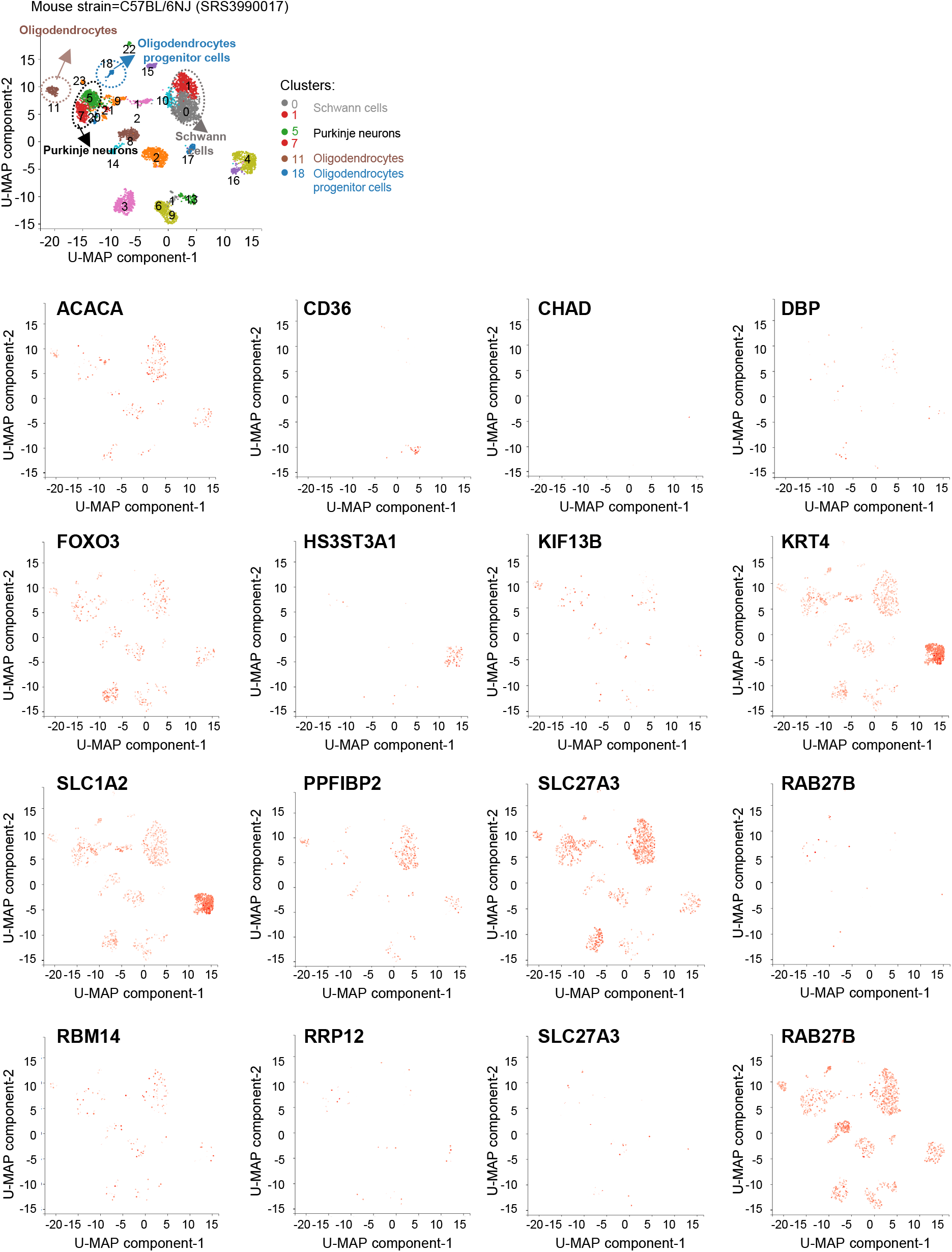
UMAP dimensional reduction visualizations of 5240 cells from olfactory bulb (mouse strain=C57BL/6NJ). Color represents normalized log expression of selected genes. Each point represents a cell. Original data were obtained from Gene Expression Omnibus (GEO; GSM3449587) and Sequencing data was handled using PanglaoDB (https://panglaodb.se/).

**Figure S6.**
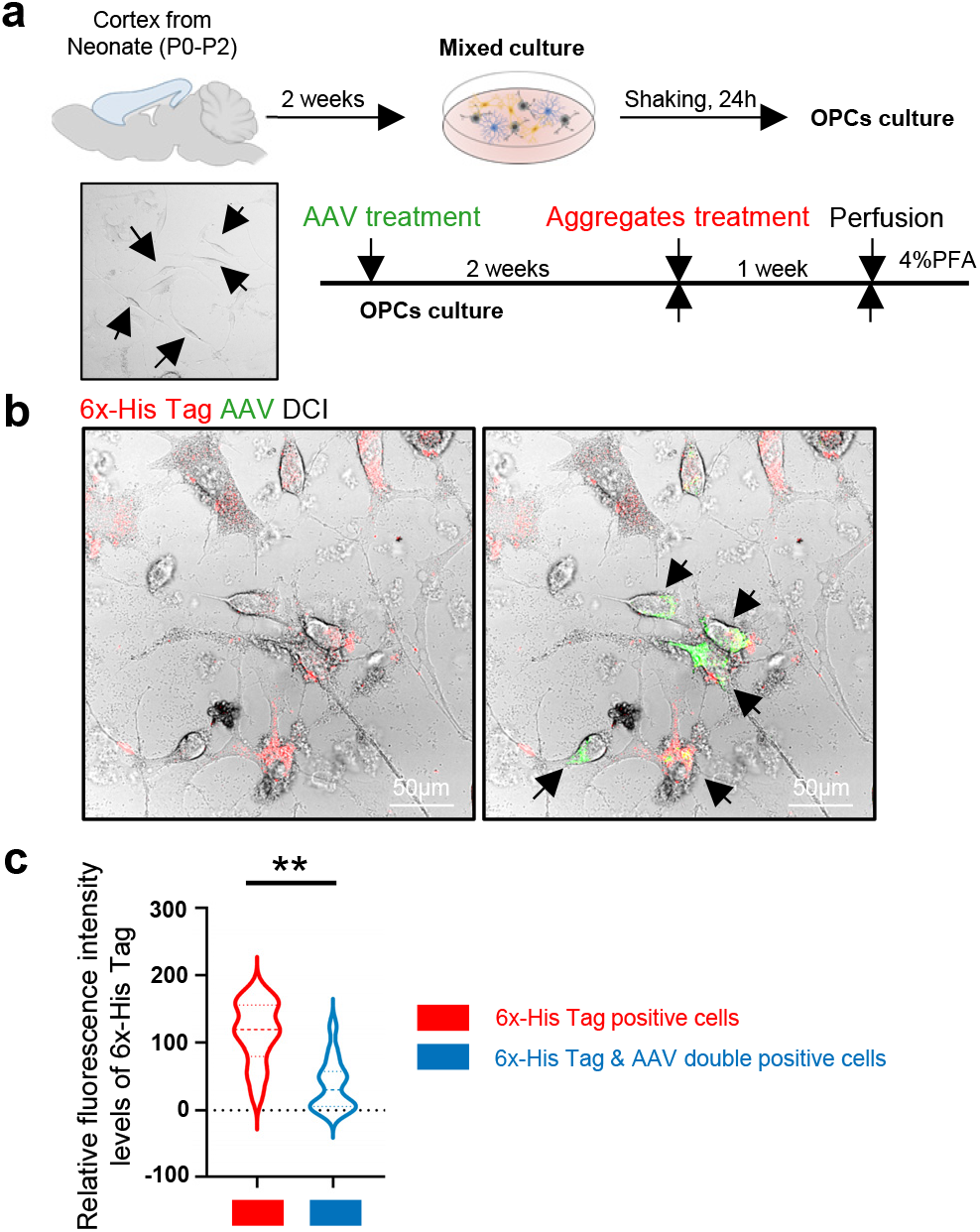
Downregulation of epsin2 decreased hαSyn/His-hFABP7 aggregates uptake in OPCs. A, Schematic diagram and treatment procedure of OPCs from the cortex of P1-2 mouse pups. B, Confocal images show aggregates in red, AAV in green. C, Quantification of B, suggested that the levels of hαSyn/His-hFABP7 aggregates uptake in OPCs decreased by epsin2 knock down. The data are presented as the mean ± the standard error of the mean, were obtained using Unpaired Student’s t-test. **P < 0.01. αSyn, alpha-synuclein.

**Figure S7:**
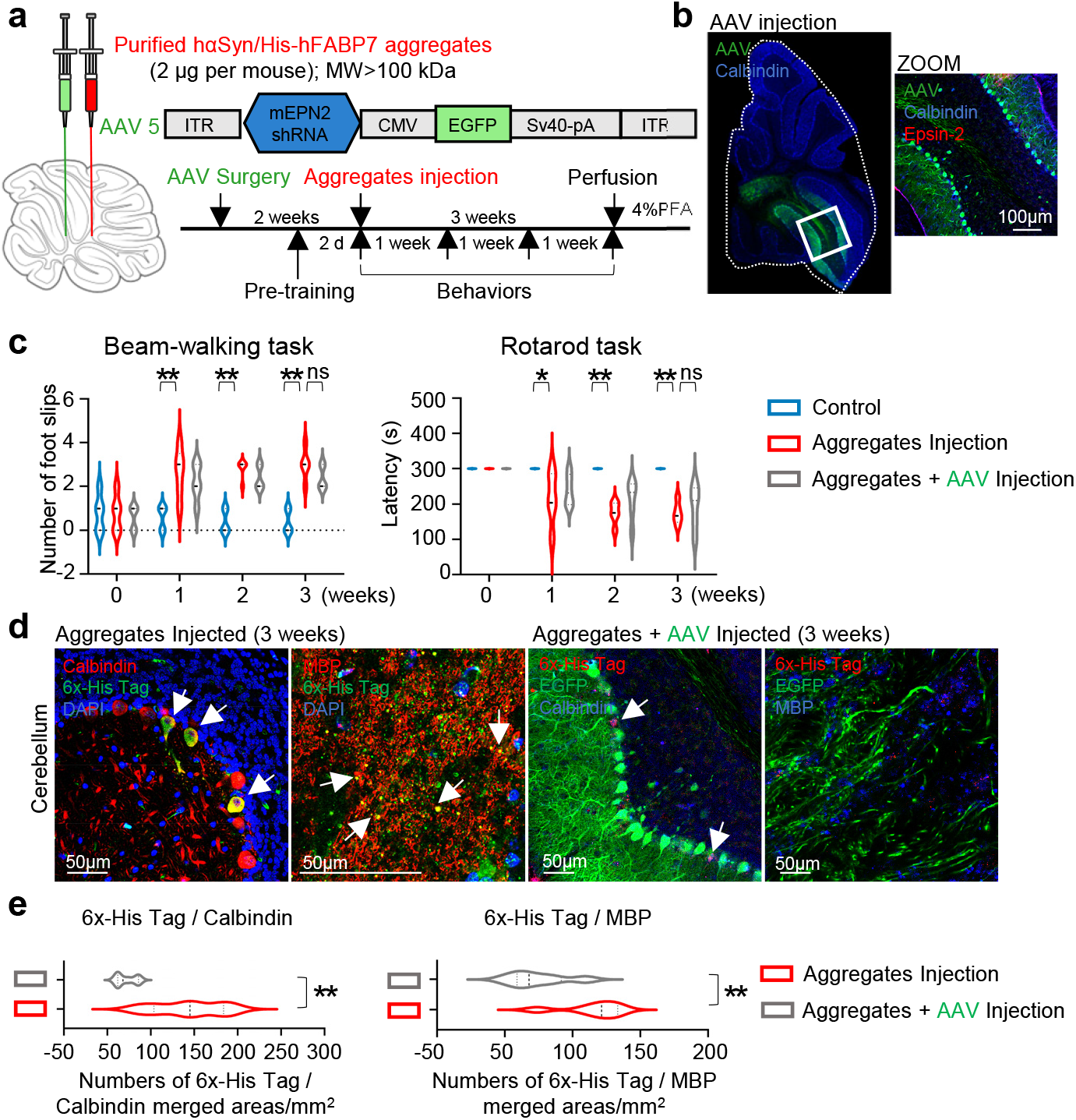
Knock down of epsin-2 decreased FABP7/α-syn hetero-aggregates uptake. a, Scheme of aggregates, and AAV injection and experimental protocol. b, Confocal images show distribution of AAV (green), epsin-2 (red), and DAPI (blue) in cerebellum. c, Behavioral tasks were carried out at beginning and after aggregates or AAV injection for 1 week, 2 weeks and 3 weeks. d, Confocal images show the propagation of injected aggregates in cerebellum. e, Quantification of d, suggested decreased levels of hαSyn/His-hFABP7 aggregates uptake in purkinje neurons and oligodendrocytes in AAV injected mice. The data are presented as the mean ± the standard error of the mean, were obtained using student’s t-test and two-way analysis of variance *P < 0.05 and **P < 0.01. αSyn, alpha-synuclein; FABP7, fatty acid binding protein 7; MBP, myelin basic protein; NS, nonsignificant.

**Table S1.** Raw data for Go and KEGG pathway analysis in Figure 1B.

| Category | Term | Count | % | PValue | Pop Hits | Pop Total | Fold Enrichment |
| --- | --- | --- | --- | --- | --- | --- | --- |
| KEGG | Olfactory transduction | 120 | 4.28878 | 2.64E-08 | 1167 | 8977 | 1.628015578 |
| KEGG | Pancreatic secretion | 22 | 0.78628 | 7.29E-06 | 114 | 8977 | 3.055385377 |
| KEGG | Salivary secretion | 16 | 0.57184 | 2.40E-04 | 85 | 8977 | 2.980226165 |
| KEGG | Yersinia infection | 19 | 0.67906 | 0.00181 | 134 | 8977 | 2.244899839 |
| KEGG | Neuroactive ligand-receptor interaction | 37 | 1.32237 | 0.01135 | 386 | 8977 | 1.517618408 |
| KEGG | Retrograde endocannabinoid signaling | 18 | 0.64332 | 0.01169 | 148 | 8977 | 1.925568426 |
| KEGG | Relaxin signaling pathway | 16 | 0.57184 | 0.01554 | 129 | 8977 | 1.963714915 |
| KEGG | Human cytomegalovirus infection | 25 | 0.8935 | 0.03255 | 256 | 8977 | 1.546137842 |
| KEGG | Spinocerebellar ataxia | 16 | 0.57184 | 0.03195 | 141 | 8977 | 1.796590241 |
| KEGG | Toll-like receptor signaling pathway | 13 | 0.46462 | 0.02297 | 100 | 8977 | 2.058218695 |

**Table S2.**
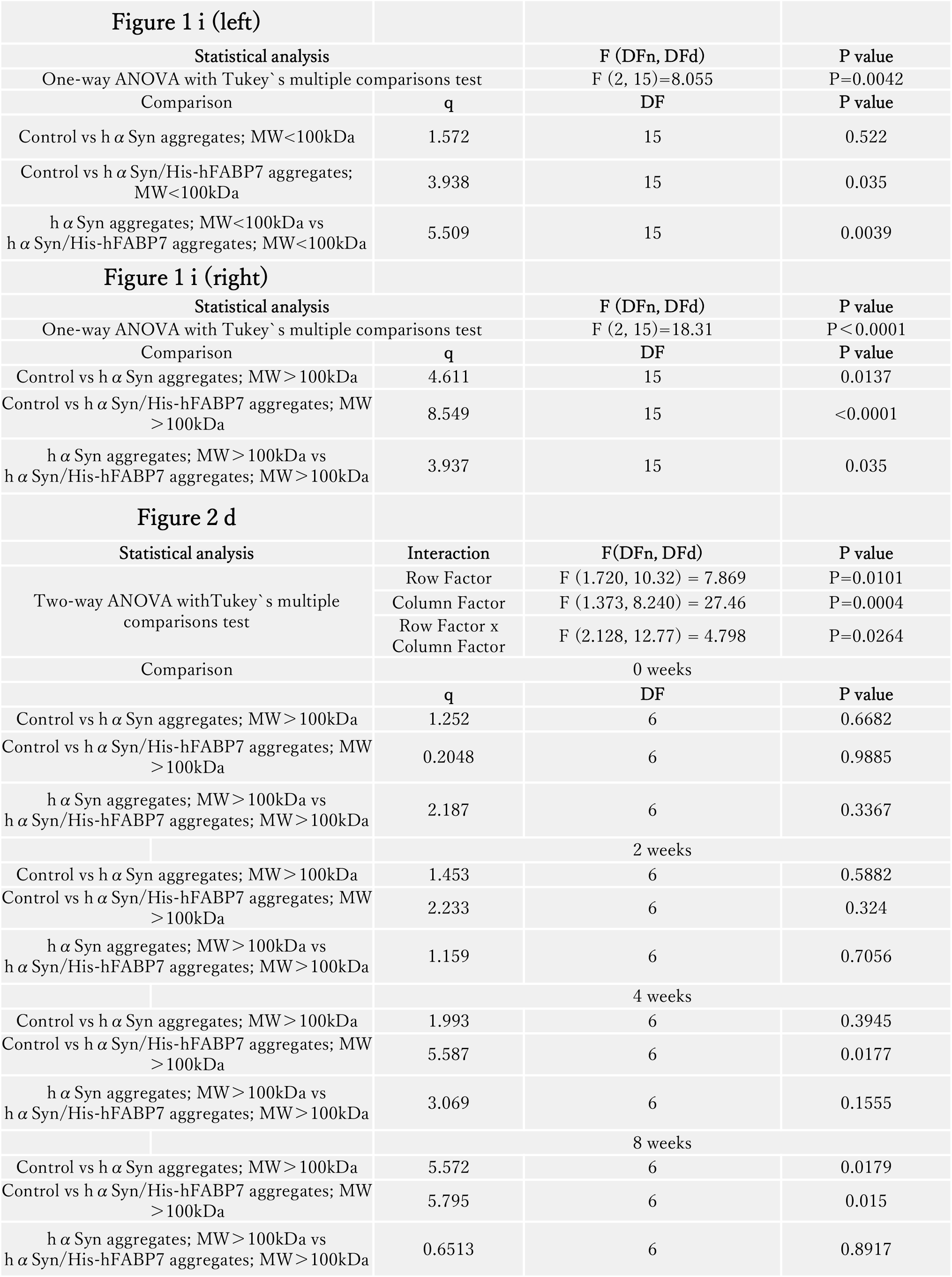
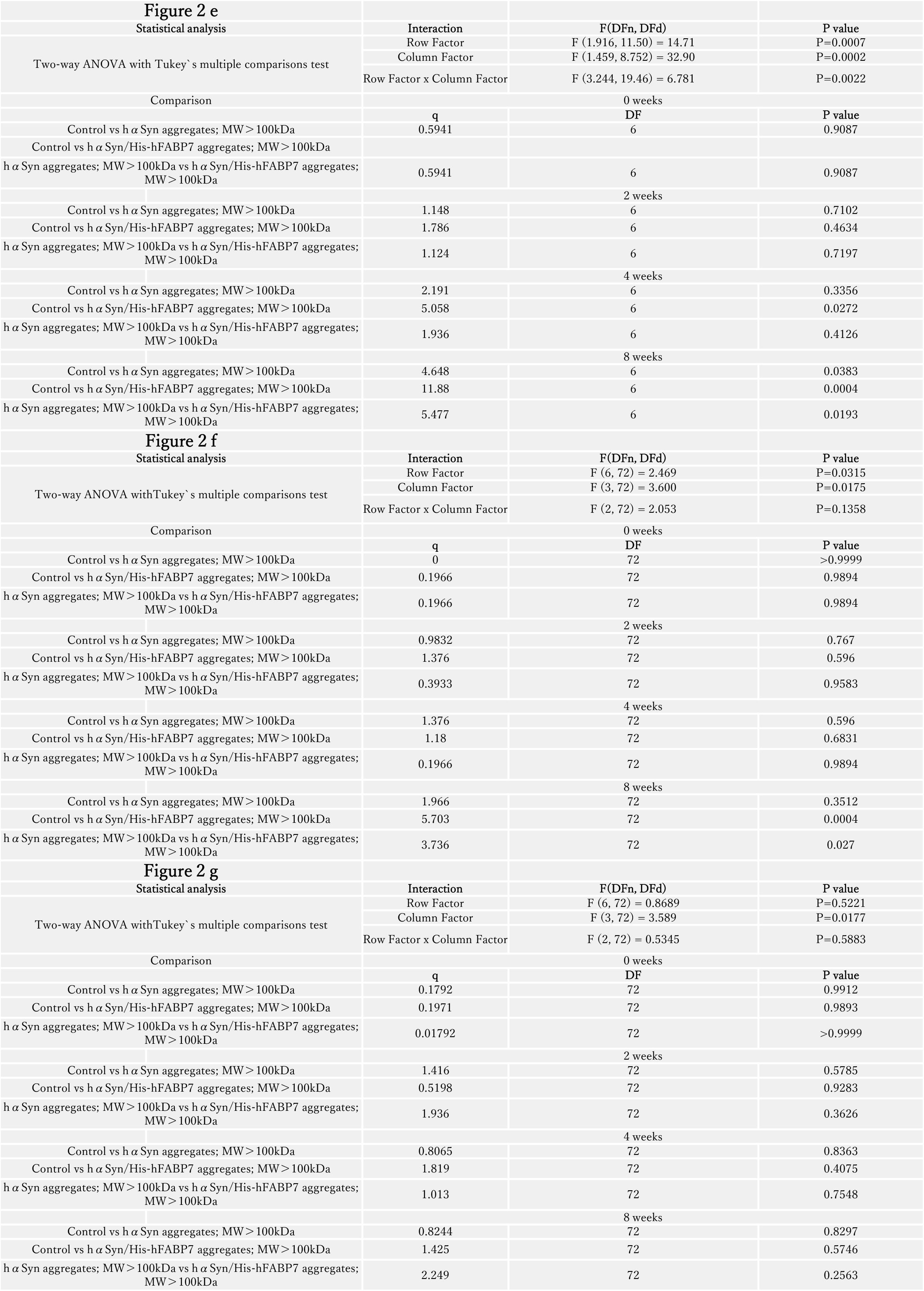
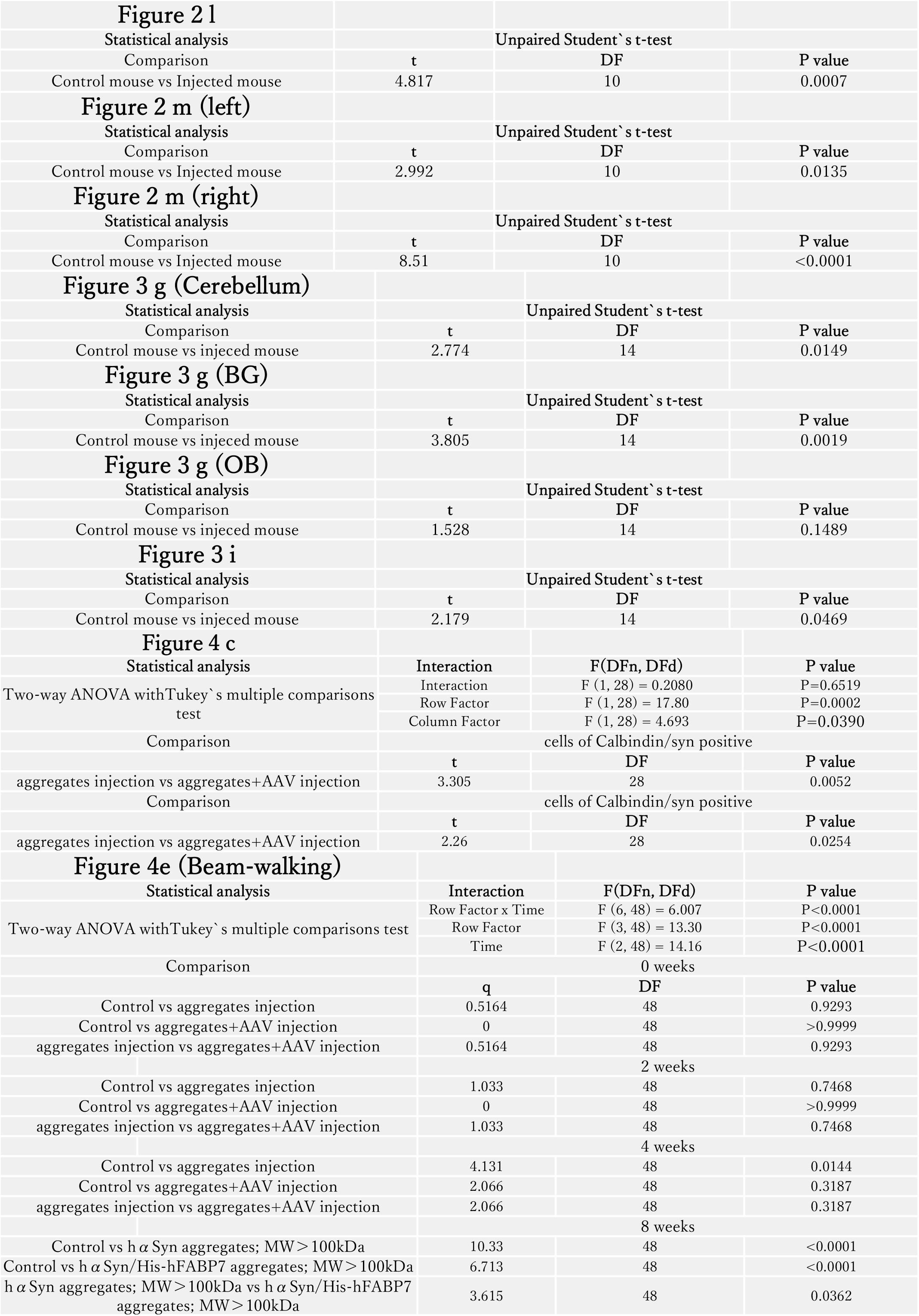
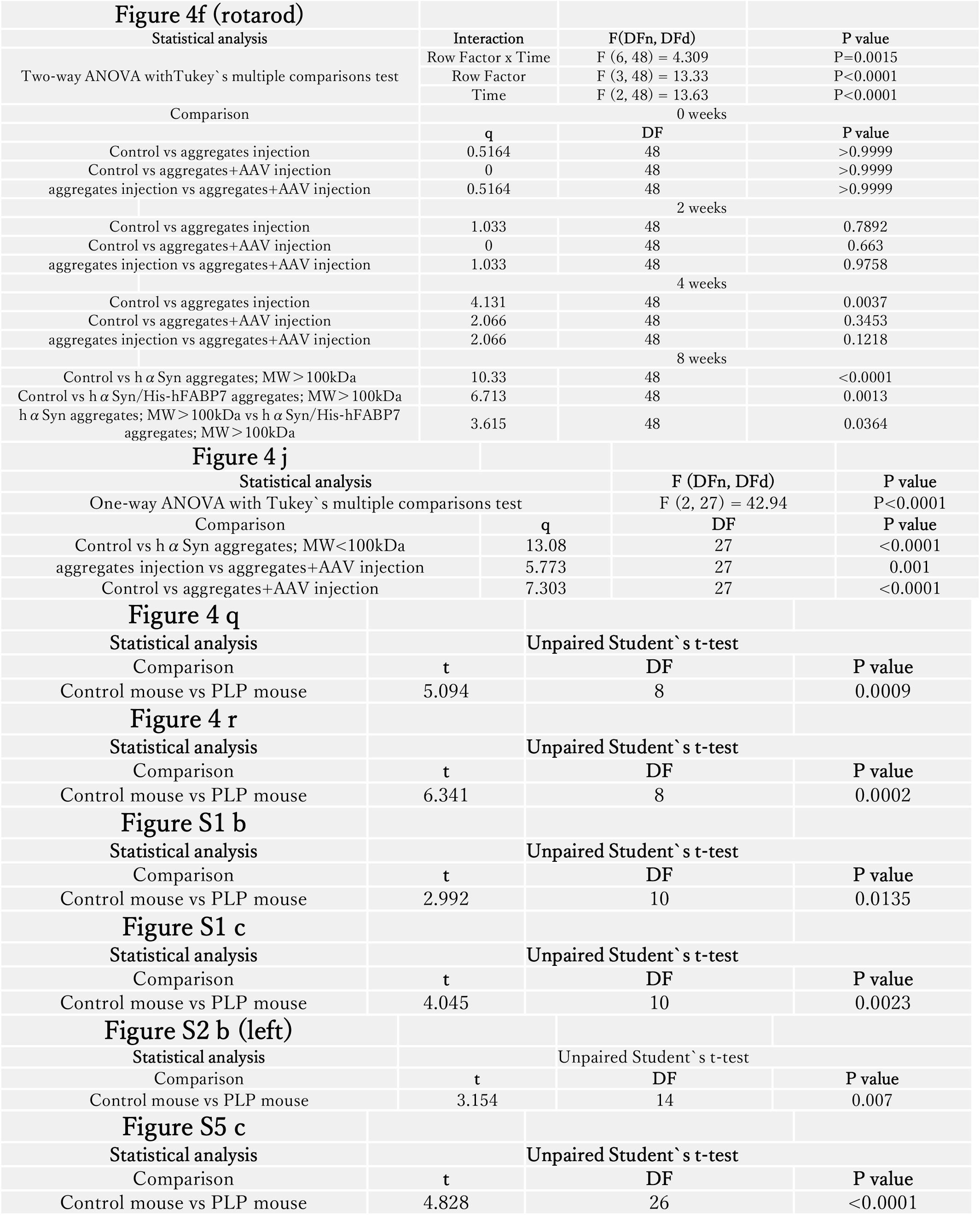
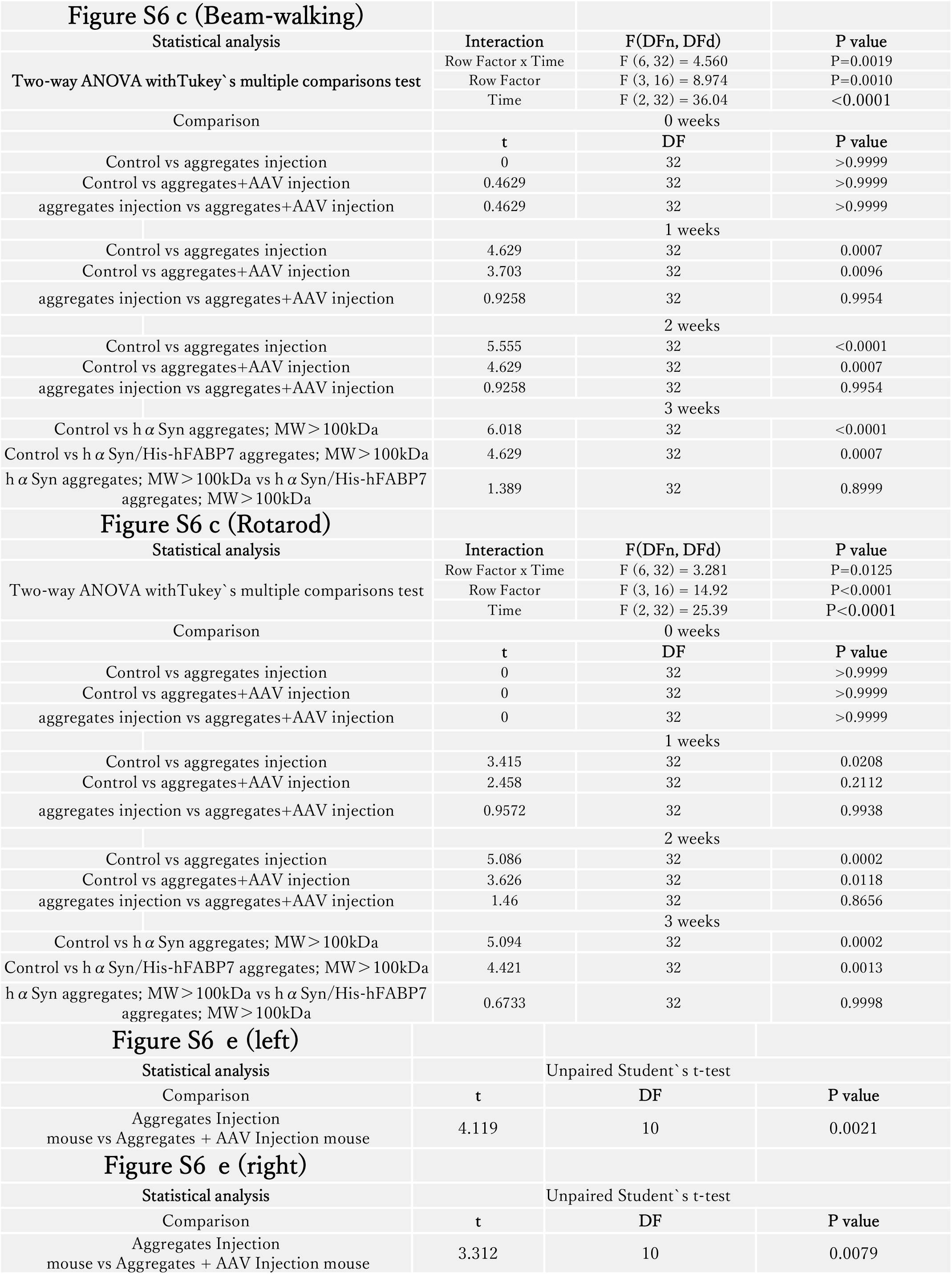
Extended Data.

### Full-size scans of western blots

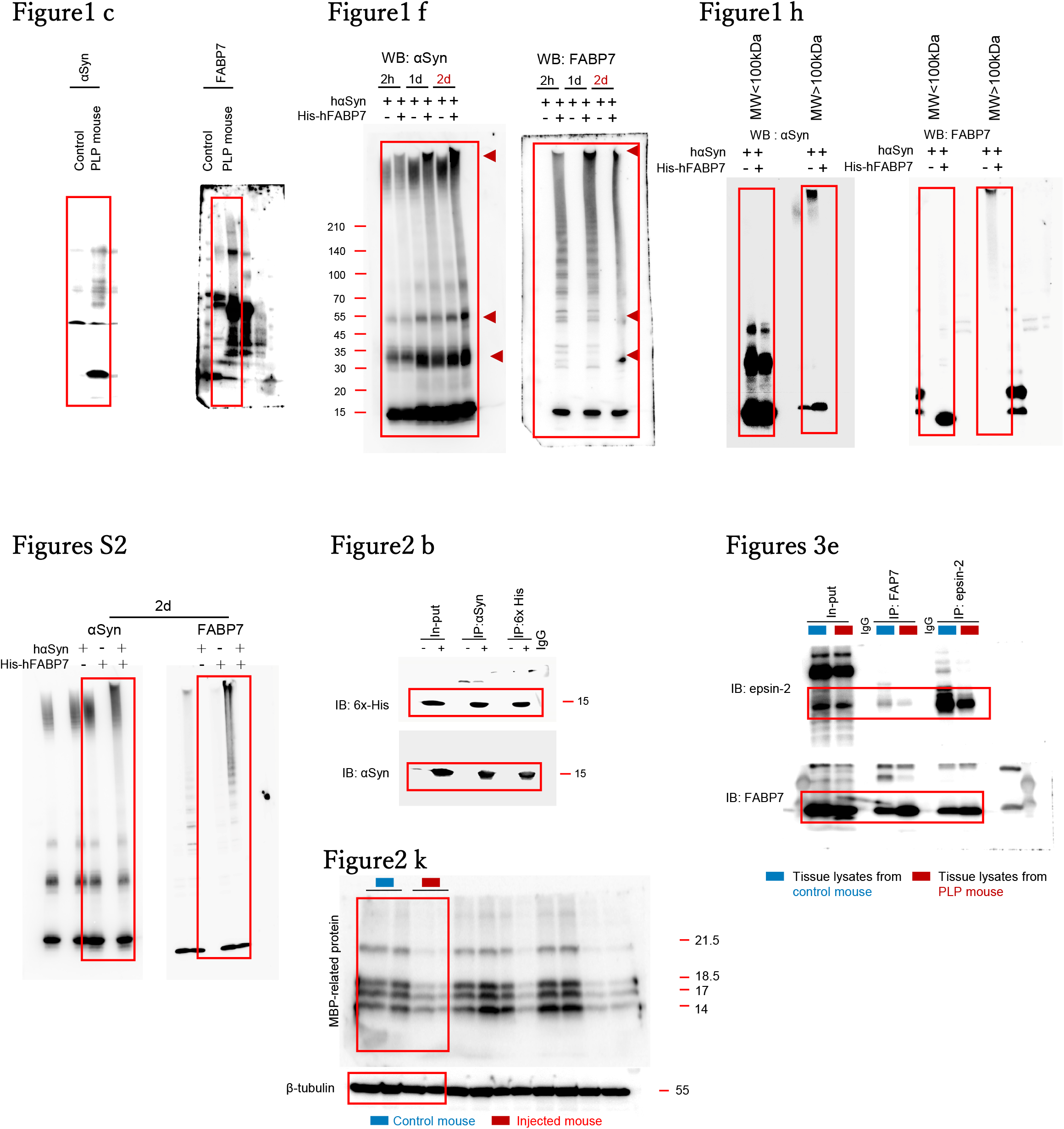

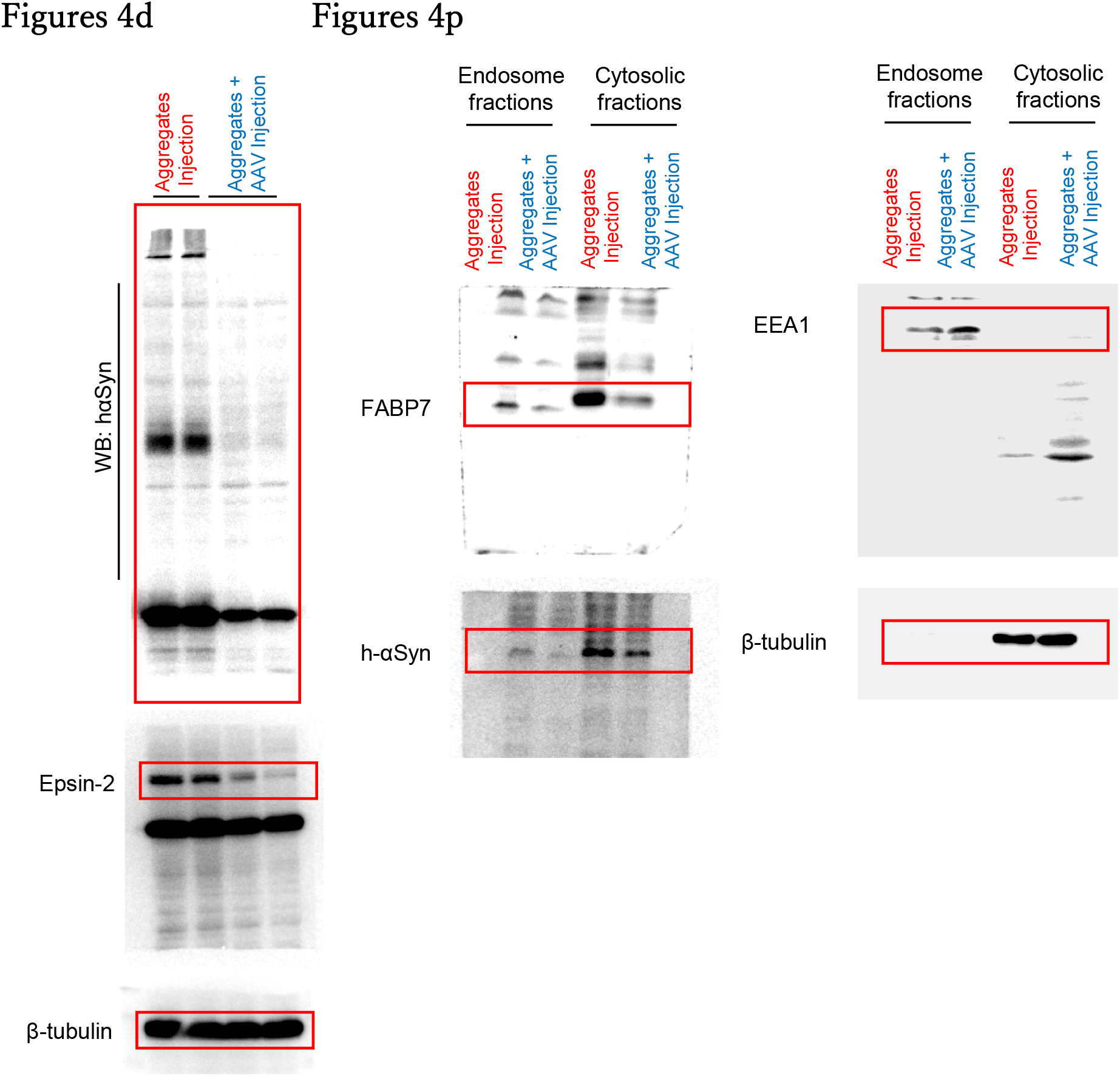

